# DetectGxT: detecting gene-by-treatment interactions on molecular count phenotypes accounting for allelic additivity

**DOI:** 10.64898/2026.07.30.740713

**Authors:** Yuriko Harigaya, Michael I. Love, William Valdar

## Abstract

**Motivation:** Identifying the mechanisms by which genetic variants affect the molecular response to an applied treatment is important across multiple biological fields, and an effective approach to this end is interaction molecular QTL mapping. However, the statistical models commonly used to detect such gene-by-treatment interactions (G×T) are non-trivially misspecified, and this can lead to decreased power.

**Results:** We developed an R software package, DetectGxT, that uses nonlinear regression to more accurately model the relationship between the genotype and the transformed molecular count phenotypes. It also optionally models donor or polygenic random effects. Simulations show that nonlinear regression can increase the power to detect interactions. In existing interaction expression QTL mapping data from primary human neural progenitor cells, nonlinear and linear regression approaches identified overlapping but distinct sets of gene-SNP pairs with significant G×T interactions. Overall, our results suggest an advantage of nonlinear regression over linear regression in detecting G×T interactions on molecular phenotypes.

**Availability:** The DetectGxT software is available at https://github.com/yharigaya/detectgxt.

## Introduction

Gene-by-treatment interaction (G×T) describes when the response of an individual to an applied treatment differs according to their genetics. To the extent that an applied treatment resembles an environment, G × T is also a type of gene-by-environment interaction, or G × E (Herrera-Luis et al., 2024; Motsinger-Reif et al., 2024). Identifying and understanding G×T is key to advancing personalized medicine, as well as other fields such as toxicology and agriculture. In particular, to better understand biological mechanisms underlying G×T, it is valuable to identify G×T at the molecular level, where responses comprise count-based molecular phenotypes, such as gene expression or chromatin accessibility, associated with genomic features such as genes or ATAC-seq peaks.

A natural approach for identifying molecular G × T is molecular QTL mapping on biological samples from genetically distinct individuals under two or more treatment conditions. Such studies are well-suited to experiments on model organisms (e.g., Mosedale et al. 2017) but, at least historically, have had limited success in humans due to the need for careful matching of genetics and environment between treatment and control. Recently, however, multiple studies have utilized in vitro cell systems derived from genetically diverse human donors, where analysis can be performed under physiological or clinically relevant stimuli. These studies used datasets containing an equal number of samples in the control and treated conditions for a given genomewide genotype, which we call “paired” data, and undertook various analytic approaches, which can be categorized into three groups:

1. Fold change methods: use the log fold change, e.g., in gene expression, between treated and control as the quantitative trait to be mapped (Barreiro et al., 2012; Çalışkan et al., 2015; Nédélec et al., 2016; Mosedale et al., 2017).
2. One-step interaction QTL mapping: jointly model log-transformed count data in the control and treated conditions using a linear model that includes genotype, treatment, and G×T interaction terms. Test for the significance of the G × T term (or the genotype and G × T terms) at all candidate *cis*-regulatory SNPs in the genome (Knowles et al., 2018).
3. Two-step interaction QTL mapping: first, perform condition-stratified QTL mapping—mapping QTL in the control and treated conditions separately using (typically) a standard linear model applied to log-transformed count data; second, apply the interaction model above to the combined data, testing only the union set of feature-SNP pairs from the two conditions (Alasoo et al., 2018; Matoba et al., 2024).

This work focuses on the last of these, two-step interaction QTL mapping. Notably, interaction QTL mapping, whether one- or two-step, does not—unlike fold change methods—require samples to be paired, and has been successfully applied to population-level, unpaired tissue samples to study gene-by-environment interaction on gene expression (Kim-Hellmuth et al., 2020; Oliva et al., 2020; Kasela et al., 2024); this unpaired setting is therefore also considered here.

Although the aforementioned studies identified many significant interactions, the statistical model underlying the QTL mapping approach may have been suboptimal: molecular counts were processed with a variance-stabilizing transformation, such as the log transformation, and the relationship between the genotype and the transformed phenotype was assumed to be linear. We and others, however, have found that the linearity typically holds on the original count scale, not on the transformed scale (Sun, 2012; Mohammadi et al., 2017; Palowitch et al., 2018; Harigaya et al., 2025), consistent with the “allelic additivity” assumption, which posits that the total molecular count in a diploid cell is the sum of molecular counts from the two alleles on the original count scale (Sun, 2012). This assumption can be satisfied by a nonlinear model (Mohammadi et al., 2017; Palowitch et al., 2018) (Supplementary Text S1.1), although at greater computational expense that may not always be pragmatic for detecting marginal genotype effects in large-scale analyses. In more focused analyses, though, such as in the second step of two-step interaction mapping, computational demands are less and nonlinear modeling could be practical. We have previously observed that, in the related task of G×T classification, nonlinear regression outperforms commonly used linear regression approaches (Harigaya et al., 2025). Here we ask if the same is true for G × T detection, specifically, is the linear model assumption sufficient for detection of G×T interactions? And more generally, what modeling considerations are important for power to detect interactions?

Another critical aspect of molecular QTL mapping is the use of mixed-effects models, where error correlation between samples from the same donor or population structure among donors is explicitly modeled using random effects. Accounting for correlation between samples due to repeated measurements of the same experimental units, such as donors, is essential to reduce false positives in differential gene expression analysis (Nguyen and Nettleton, 2020), and including polygenic random effects to control for polygenicity and population structure helps reduce spurious molecular QTLs (Lee, 2018).

Given these considerations, it is desirable to perform molecular interaction QTL mapping using nonlinear mixed-effects models to account for the allelic additivity and within-donor correlations. However, such analysis cannot readily be applied using existing computational tools (Supplementary Text S1.2). Herein, we provide a software package, DetectGxT, that allows users to perform interaction molecular QTL mapping accounting for allelic additivity, and optionally including donor or polygenic random effects. In the context of two-step interaction QTL mapping, our simulation experiments show that nonlinear regression in the second step increases the power to detect interactions as compared with linear regression. We also apply DetectGxT to existing interaction expression QTL (eQTL) mapping data from primary human neural progenitor cells (hNPCs) (Matoba et al., 2024) and obtain overlapping but distinct sets of gene-SNP pairs with significant G×T interactions from nonlinear and linear regression approaches. These results suggest that the use of nonlinear regression rather than linear regression can be advantageous in detecting G*×*T interactions.

### Software description

DetectGxT identifies significant G×T interactions on molecular count phenotypes, such as gene expression (RNA-seq) and chromatin accessibility (ATAC-seq). This software is primarily designed for the second step in two-step interaction QTL mapping: interaction mapping of candidate feature-SNP pairs with significant genotype effects in at least one condition.

The software vignette explains the entire data analysis procedure. In the first step (Fig. 1A), condition-stratified molecular QTL mapping is performed using standard software packages, such as MatrixEQTL, TensorQTL, and limix_qtl (Shabalin, 2012; Taylor-Weiner et al., 2019; Lippert et al., 2014). These assume a linear relationship between the genotype and transformed molecular phenotypes: although this can lead to inaccurate effect estimation, this is unlikely to be a major issue for detection in a single condition, and is highly computationally efficient. SNPs with significant associations can be filtered using standard approaches, including choosing pergene lead SNPs, LD clumping, conditional analysis, or fine-mapping (Purcell et al., 2007; Genetic Investigation of ANthropometric Traits (GIANT) Consortium et al., 2012; Wang et al., 2020). After this step, we select the union set of feature-SNP pairs with significant association in either condition.

**Figure 1.**
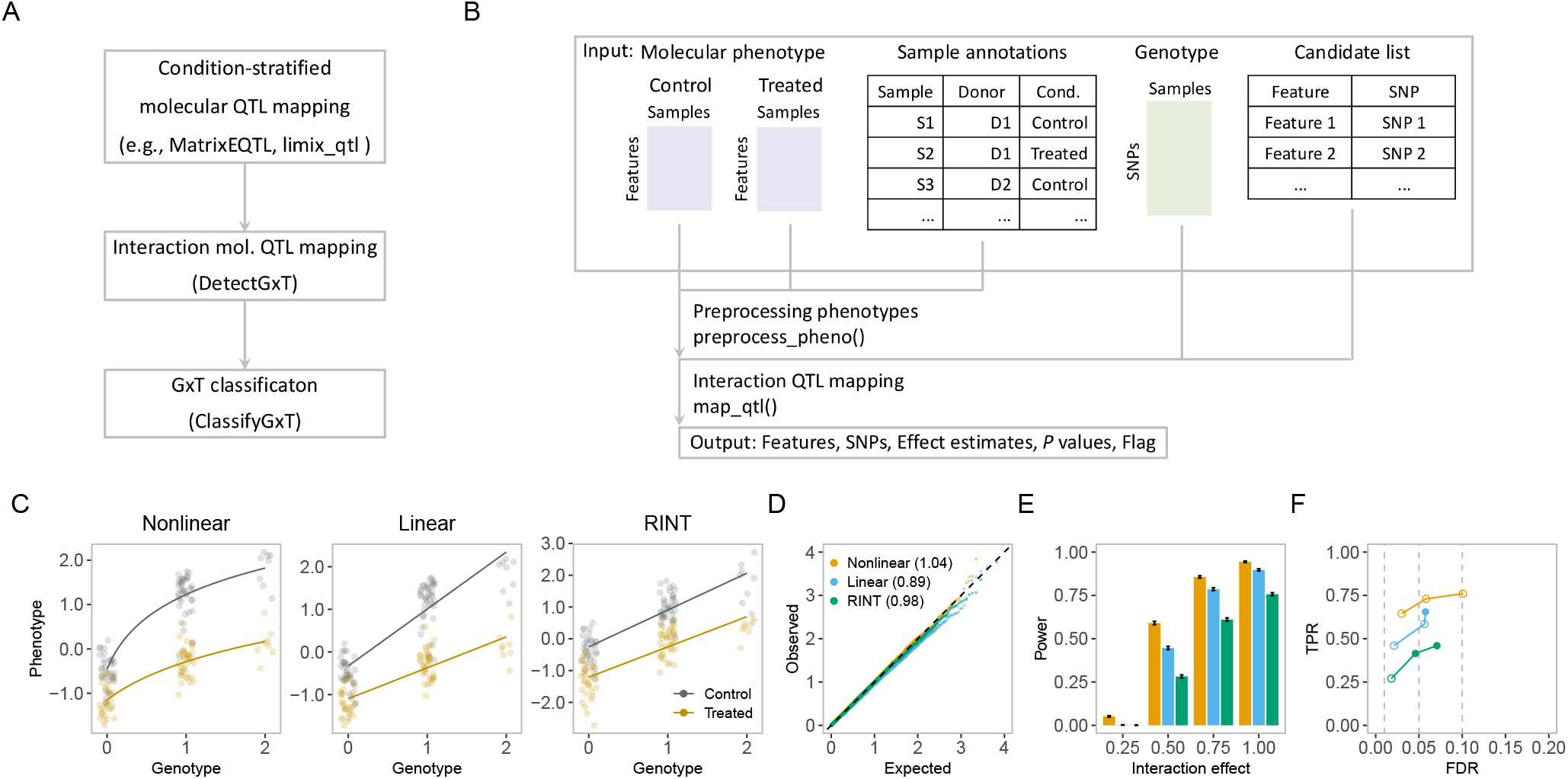
Overview of DetectGxT. (A) Overall workflow. (B) DetectGxT provides the `map_qtl()` function to test for G*×*T interactions accounting for allelic additivity. Inputs are molecular count data in control and treated conditions, sample annotations, genotype data, and a list of candidate feature-SNP pairs. Outputs are effect estimates, *P* values, and flags for low frequency of either homozygote. Optional data preprocessing with `preprocess_pheno()`. (C) Example molecular count data in control and treated conditions: The y-axis is on the log scale (left, middle), or RINT-transformed (right). Fitted curves from nonlinear regression (left), and linear regression (middle, right). (D–F) Results for simulation with no random effects in the paired setting (n = 156). (D) Calibration of *P* values under the null (no G*×*T or random effect): Q-Q plot with theoretical (x-axis) and empirical (y-axis) values of − log_10_(*P*); in parentheses are genomic inflation factors (*≈* 1 if well-calibrated).(E) Power to detect G*×*T interaction under varying G*×*T effect sizes, *β_g×t_*. (F) Overall performance assessed by the empirical true positive rate (TPR) vs empirical false discovery rate (FDR) at the nominal FDR of 0.01, 0.05, and 0.1. Closed circles indicate empirical FDR *<* nominal FDR; open circles, the opposite.

In the second step, for each of the feature-SNP pairs selected above, we use the nonlinear interaction model to analyze their molecular count data in control and treated conditions jointly (Fig. 1B and Supplementary Text S2). Prior to this step, the input molecular phenotype data should be preprocessed appropriately, as in previous studies (Alasoo et al., 2018; Matoba et al., 2024) or using our function `preprocess_pheno()` (Supplementary Text S2.3). Also prior to this step, as a conservative measure, we recommend excluding “AA-AB” SNPs, where reference or alternative allele homozygotes are missing in at least one condition, because they can create practical non-identifiability and misestimation of genotype and interaction effects in the nonlinear model; the excluded SNPs likely correspond to a small fraction in typical datasets (Supplementary Text S3.3). Feature-SNP pairs with significant G×T interactions can then be identified using the `map_qtl()` function (Fig. 1B). Subsequently, users can optionally group the interaction molecular QTLs with significant G ×T interactions into different types using ClassifyGxT (Harigaya et al., 2025) (Fig. 1A).

## Results

We first evaluated the G×T detection procedure using simulations. Simulated data were generated using nonlinear models based on the allelic additivity assumption supported by previous studies (Mohammadi et al., 2017; Palowitch et al., 2018; Harigaya et al., 2025). To these data, we applied three methods: nonlinear (the data-generating model); linear (linear additive model, assuming linearity on the transformed scale); RINT (linear after rank-based inverse normal transformation Beasley et al. 2009) (Fig. 1C).

We considered two settings. In the “paired” setting, the data contain one sample per combination of genotype and condition, as in the previous study of an in vitro cell system (hNPCs with growth stimulation, Matoba et al. 2024). The data comprised one observation per condition for 78 or 156 donors, corresponding to total sample sizes of 156 or 312, respectively. Data were generated with a donor random effect, as within-donor correlation is a strong feature of this setting. Polygenic random effects were not simulated or modeled in this case: doing so would require a second random effect (although see Supplementary Text S4), and polygenicity and stratification can be accounted for by regressing out genotype principal components (Kim-Hellmuth et al., 2020).

In the “unpaired” setting, the data contain one sample from each donor, with some donors in the baseline condition and others in the treated condition. Total sample sizes were 156 and 312. This type of setting includes gene-by-sex analyses of molecular phenotype data in genotyped individuals from consortia, such as GTEx and TOPMed (Oliva et al., 2020; Kasela et al., 2024). Data were generated with a polygenic random effect to account for possible population structure. To account for different types of population structure, simulations were repeated using genomic relationship matrices (GRMs) derived from each of four ancestry populations (The 1000 Genomes Project Consortium et al., 2015) (Supplementary Text S2.6.3).

In both settings, we simulated six scenarios by generating data with three levels of random effect standard deviation (zero, medium, and high), performing analyses with or without a random effect term in the model (see Supplementary Table S1), and including only SNPs for which both homozygotes are observed in both conditions (see Supplementary Text S3.3 and Figs. S1–S14). Models were evaluated using four metrics: calibration of *P* values (defined here as uniformity of *P* values under the null), statistical power, overall performance via true positive rate (TPR) and false discovery rate (FDR), and quality of effect estimation (Supplementary Text S2.5). We note that DetectGxT showed adequate computational efficiency across all scenarios and replicates (Figs. S15 and S16).

Analyses of the paired data revealed a clear advantage for the nonlinear approach, with calibration and power both depending critically on whether a donor random effect was included. Specifically, when modeling with a random effect term, *P* values were generally well-calibrated across models (except for RINT being anti-conservative at the larger sample size; Figs. 1D and S17), but the nonlinear model consistently outperformed the other methods in terms of power and overall performance (Figs. 1E, 1F, S18, and S19), yielding effect estimates that largely matched true values, as compared with the other methods, which showed substantial estimation bias (Figs. S20–S25A). When the data were generated with random effects but modeled without a random effect term, all metrics were degraded regardless of the functional relationship, suggesting the importance of accounting for donor random effects (Fig. S17).

Analyses of the unpaired data revealed that the nonlinear approach maintained calibration and power over linear and RINT approaches. Specifically, *P* values were well-calibrated for the nonlinear approach, regardless of whether a polygenic term was included (Figs. S26–S29), but were skewed toward zero (anti-conservative) for linear and RINT approaches (Figs. S26–S29). The lack of impact of the polygenic random effect term is likely due to the randomly drawn genotypes, which suffice for the comparison of the modeling approaches. We would expect to observe better calibration for mixed-effects models when using real genotypes. The nonlinear approach yielded the largest power across G × T effect sizes (Figs. S30–S33), the best overall performance (Figs. S34–S37), and superior estimation as compared with the linear and RINT approaches (Figs. S38–S43A), similarly to the results for the paired data. These observations were consistent across ancestry populations, suggesting the general superiority of the nonlinear approach.

We applied this method to a previously published RNA-seq dataset generated in hNPCs derived from 78 genetically diverse fetal donors, with and without growth stimulation (Matoba et al., 2024). The data are completely paired, and we included the polygenic random effect in the model to allow for direct comparison between the previous and current results. With linear mixed-effects models, we observed a high degree of concordance between the studies. Specifically, our implementation of linear mixed-effects models identified 79 out of 83 previously identified interaction eQTLs. We also observed that a large portion of interaction eQTLs were shared between the nonlinear and linear approaches, whereas a handful of gene-SNP pairs were unique to each approach (Fig. S44).

## Conclusion

DetectGxT enables interaction molecular QTL mapping using nonlinear regression while accommodating donor or polygenic random effects. The method improves power in our simulations and may facilitate reanalysis of existing molecular QTL datasets.

## Data availability

The detectgxt R package is publicly available on GitHub (https://github.com/yharigaya/detectgxt) and archived on Zenodo (DOI: 10.5281/zenodo.21460613). The software vignette can be viewed at https://yharigaya.github.io/detectgxt/. All code to generate the results in this paper can be found on GitHub (https://github.com/yharigaya/detectgxt-paper) and Zenodo (DOI: 10.5281/zenodo.21446264). The experimental data used in this study were published in Matoba et al. (2024) (https://doi.org/10.1038/s41593-024-01773-6), with code used for that paper available on Bitbucket (https://bitbucket.org/steinlabunc/wnt-rqtls/). The data can be accessed via the Database of Genotypes and Phenotypes (dbGaP) at https://www.ncbi.nlm.nih.gov/gap/ with the accession number phs003642.v1.p1.

## Acknowledgments

Portions of code and documentation were generated with the assistance of Claude Code (Anthropic) and Codex (OpenAI). We also used Claude (Anthropic) and ChatGPT-5 (OpenAI) to improve written text. We reviewed and verified the materials produced by these tools.

