## Supplementary Text for "DetectGxT: detecting gene-by-treatment interactions on molecular count phenotypes accounting for allelic additivity"

July 23, 2026

**4 Supplementary Discussion** **12**

**5 Supplementary Tables** **15**

**6 Supplementary Figures** **17**

### 1 Supplementary Notes

#### 1.1 Review of allelic additivity

In this section, we briefly describe a statistical model for performing molecular QTL mapping under the allelic additivity assumption in a single condition, which represents a foundation for our molecular interaction QTL mapping method (Section 2.2). The model was originally proposed by two independent studies [1, 2]. We use our own notation that differs from that of the previous studies to ensure consistency with our extended model for G×T analysis. The model is cast as

$$\begin{aligned} y_i &= \log(\mu_g(g_i)) + \varepsilon_i, \\ \mu_g(g_i) &= (1 - \tfrac{g_i}{2})\exp(\beta_0) + (\tfrac{g_i}{2})\exp(\beta_0 + 2\beta_g), \\ \varepsilon_i &\overset{\text{iid}}{\sim} \mathcal{N}(0, \sigma^2), \end{aligned}$$

where $i$ indexes samples, $y_i$ is the log-transformed count phenotype, $\mu_g(\cdot)$ is a nonlinear function, $g_i$ is the genotype coded as $\{0, 1, 2\}$ or the imputation-based allelic dosage, $g_i \in [0, 2]$, $\varepsilon_i$ is the residual error with the variance $\sigma^2$, and $\beta_0$ is the intercept. The parameter $\beta_g$ represents half the difference in the expected values of $y_i$ for the reference allele homozygote ($g = 0$) vs the alternative allele homozygote ($g = 2$). We note that

$$\begin{aligned} \mu_g(g = 0) &= \exp(\beta_0), \\ \mu_g(g = 2) &= \exp(\beta_0 + 2\beta_g), \\ \mu_g(g = 1) &= \tfrac{1}{2}\mu_g(g = 0) + \tfrac{1}{2}\mu_g(g = 2), \end{aligned}$$

which illustrates the linearity between the genotype and the phenotype in the original count scale.

#### 1.2 Limitations of existing R packages for nonlinear mixed-effects modeling

Although the lme4 and nlme R packages can be used to fit nonlinear mixed-effects models with donor random effects, it is not intuitive for users to specify custom functions to accomplish this analysis [3, 4]. Another limitation is that the packages do not provide an option to include a custom covariance matrix, such as a genetic relatedness (kinship) matrix to model polygenic random effects. Although the lme4qtl package provides the option to model polygenic effects, it only allows for a linear relationship between the genotype and the phenotype [5].

### 2 Supplementary Methods

We note that portions of Sections 2.1, 2.2, 2.3, and 2.6 have been adapted from our previous manuscript [6].

#### 2.1 Dataset and preprocessing

The experimental data in primary human neural progenitor cells (hNPCs) were generated in a previous study [7]. The data consisted of genotypes and molecular count data representing gene expression, derived from 78 genetically diverse donors with and without growth stimulation. For each gene and donor, the data contained expression measurements in control samples treated with vehicle and those treated with CHIR, an activator of the Wnt pathway, summing to 156 observations. Among 3,401 autosomal gene-SNP pairs that exhibited significant associations in at least one of the control and treated conditions, we focused on 3,083 pairs, excluding the AA-AB SNPs (i.e., cases where reference or alternative allele homozygotes are missing in at least one of the control and treated conditions). Processed gene expression data were obtained from the previous study [7]. Rank-based inverse normal transformation (RINT) [8] was performed essentially as described previously [6].

#### 2.2 Modeling the relationship between genotype and molecular count phenotype

Interaction molecular QTL mapping was performed using three modeling approaches, which we call nonlinear, linear, and RINT. In the nonlinear approach, which accounts for allelic additivity, for the $i$th sample ($i = 1, \ldots, n$), we consider the alternative model

$$y_i = f_{g,t}(g_i, t_i) + \varepsilon_i, \tag{1}$$

$$\begin{aligned} f_{g,t}(g_i, t_i) = \log \big( &(1 - \tfrac{g_i}{2})(1 - t_i)\exp(\beta_0) \\ &+ (\tfrac{g_i}{2})(1 - t_i)\exp(\beta_0 + 2\beta_g) \\ &+ (1 - \tfrac{g_i}{2})(t_i)\exp(\beta_0 + \beta_t) \\ &+ (\tfrac{g_i}{2})(t_i)\exp(\beta_0 + 2\beta_g + \beta_t + 2\beta_{g\times t})\big), \end{aligned} \tag{2}$$

where $y_i$, $g_i$, $\varepsilon_i$, $\sigma^2$, and $\beta_0$ are defined as before, $t_i$ denotes an indicator variable for a treatment, $\beta_g$ represents half the difference in the expected values of $y_i$ for the reference allele homozygote vs the alternative allele homozygote *under the control condition*, $\beta_t$ is the difference in the expected values of $y_i$ in the treated condition vs the control condition *for the reference allele homozygote*, and $\beta_{g\times t}$ allows the effect of genotypes to vary between treatment groups. In the null model, $f_{g,t}(\cdot,\cdot)$ is defined as

$$\begin{aligned} f_{g,t}(g_i, t_i) = \log \big( &(1 - \tfrac{g_i}{2})(1 - t_i)\exp(\beta_0) \\ &+ (\tfrac{g_i}{2})(1 - t_i)\exp(\beta_0 + 2\beta_g) \\ &+ (1 - \tfrac{g_i}{2})(t_i)\exp(\beta_0 + \beta_t) \\ &+ (\tfrac{g_i}{2})(t_i)\exp(\beta_0 + 2\beta_g + \beta_t)\big). \end{aligned} \tag{3}$$

The linear and RINT approaches use linear regression with the alternative and null models where $f_{g,t}(\cdot,\cdot)$ is respectively defined as

$$f_{g,t}(g_i, t_i) = \beta_0 + \beta_g g_i + \beta_t t_i + \beta_{g\times t} g_i t_i, \tag{4}$$

and

$$f_{g,t}(g_i, t_i) = \beta_0 + \beta_g g_i + \beta_t t_i. \tag{5}$$

Here, $g_i$, $t_i$, $\varepsilon_i$, $\sigma^2$, $\beta_0$, $\beta_g$, $\beta_t$, and $\beta_{g\times t}$ are defined as before. In the linear and RINT approaches, $y_i$ denotes log- and RINT-transformed molecular count data, respectively.

Note that, for ease of exposition, confounding factors are omitted in the above equations. See Section 2.3 for details on including covariates.

#### 2.3 Accounting for covariates

In this section, we describe models that include confounding factors as fixed and random effects. We let $m$ denote the number of donors, $j$ index the donors, and $n_j$ denote the number of samples in the $j$-th donor group. The total number of samples can be written as $n = \sum_{j=1}^{m} n_j$. Then, for the $i$-th sample in the $j$-th donor group ($i = 1, \ldots, n_j$, $j = 1, \ldots, m$), the alternative model can be cast as

$$\begin{aligned} y_{i,j} &= f_{g,t}(g_{i,j}, t_{i,j}) + \mathbf{x}_{i,j}^T \boldsymbol{\gamma} + u_j + \varepsilon_{i,j}, \\ \mathbf{u} &\sim \mathcal{N}_m(0, \sigma_u^2 \mathbf{K}), \\ \varepsilon_{i,j} &\overset{\text{iid}}{\sim} \mathcal{N}(0, \sigma^2), \end{aligned}$$

where $y_{i,j}$ denotes the phenotype, $f_{g,t}(\cdot,\cdot)$ is defined as in Eq. (2), $q$ is the number of fixed-effect factors, $\mathbf{x}_{i,j} \in \mathbb{R}^q$ denotes a vector of fixed-effect factors, $\boldsymbol{\gamma} \in \mathbb{R}^q$ denotes a vector of the corresponding coefficients, $\mathbf{u} \in \mathbb{R}^m$ denotes a vector of random effects, $\sigma_u$ denotes the random effect standard deviation, $\varepsilon_{i,j}$ denotes the residual error, and $\sigma^2$ denotes the residual error variance. The matrix $\mathbf{K}$ denotes a kernel matrix and can be a known kinship matrix, which represents genetic relatedness between donors. Alternatively, when the genetic relatedness is included as a fixed effect, $\mathbf{K}$ can be set to the $m$-dimensional identity matrix $\mathbf{I}_m$. In matrix form, we have

$$\mathbf{y} \sim \mathcal{N}_n(f_{g,t}(\mathbf{g}, \mathbf{t}) + \mathbf{X}\boldsymbol{\gamma} + \mathbf{Z}\mathbf{u}, \sigma^2 \mathbf{I}_n),$$

where $\mathbf{y} \in \mathbb{R}^n$ denotes the outcome vector,

$$\begin{aligned} f_{g,t}(\mathbf{g}, \mathbf{t}) = (&f_{g,t}(g_{1,1}, t_{1,1}), \ldots, f_{g,t}(g_{n_1,1}, t_{n_1,1}), \\ &\ldots, \\ &f_{g,t}(g_{1,m}, t_{1,m}), \ldots, f_{g,t}(g_{n_m,m}, t_{n_m,m}))^T \in \mathbb{R}^n \end{aligned} \tag{6}$$

denotes the mean values, $\mathbf{X} \in \mathbb{R}^{n\times q}$ denotes a matrix of fixed-effect factors, and $\mathbf{Z} \in \mathbb{R}^{n\times m}$ denotes an incidence matrix indicating the group membership of the samples. We marginalize out the random effects $\mathbf{u}$ and obtain

$$\mathbf{y} \sim \mathcal{N}_n(f_{g,t}(\mathbf{g}, \mathbf{t}) + \mathbf{X}\boldsymbol{\gamma}, \sigma_u^2 \mathbf{Z}\mathbf{K}\mathbf{Z}^T + \sigma^2 \mathbf{I}_n). \tag{7}$$

The null model is obtained by defining $f_{g,t}(\cdot,\cdot)$ as in Eq. (3).

In our G×T analysis of the gene expression from hNPCs, we included the top 10 principal components (PCs) of the molecular count data as fixed effects and the genetic relatedness (kinship) among the donors as a random effect as in the previous study [7]. The molecular count PCs were identified in two steps. First, for each feature, we combined the library size-scaled, log-transformed molecular count data in the control and treated conditions and residualized the combined data with respect to the treatment indicator variable. We did not regress out the donor or kinship random effect since these effects are unlikely to be captured in the top PCs. Second, we obtained PCs by performing principal component analysis (PCA) on the residuals. Then, residualized phenotype data, $y_i$, was obtained by regressing out the top 10 PCs identified above from the library size-scaled, log-transformed data prior to model fitting. Although it is ideal to control for the factors simultaneously with examining the associations of the phenotype with the genotype and treatment, such a model formulation can pose computational challenges due to a large number of parameters to be estimated. Therefore, we residualized the library size-scaled, log-transformed count data with respect to the fixed-effect confounding factors and fitted models including only random effects. Note that we did not consider fixed-effect confounding factors in the simulation experiments.

We implemented this procedure in the `preprocess_pheno()` function. We note that the number of PCs can be determined based on various methods, including visual inspection of a scree plot or an automatic elbow detection algorithm [9]. To facilitate this process, `preprocess_pheno()` provides an option to return PCs.

#### 2.4 Model fitting, hypothesis testing, and multiple testing correction

Given Eqs. (2), (3), (4), (5), (6), and (7), we obtained the maximum likelihood estimates of $\boldsymbol{\beta}$, $\boldsymbol{\gamma}$, $\sigma_u$, and $\sigma$ via numerical optimization. In this procedure, we reparameterized the equations as $s^2 = \sigma_u^2 + \sigma^2$ (total variance) and $h^2 = \sigma_u^2/s^2$ (heritability). We first fixed $h^2 \in [0, 1-\delta]$ and optimized $\boldsymbol{\beta}$ and $s^2$ using the BFGS algorithm in the `optim()` function from the `stats` R package [10]. We then optimized $h^2$ via Brent's method [11]. To avoid numerical instability near the boundary, we set $\delta = 1 \times 10^{-2}$. We implemented this procedure in the `map_qtl()` function.

We used the likelihood ratio test (LRT) [12] to determine the significance of G×T interactions. The likelihood ratio statistic for testing $H_0: \theta \in \Theta_0$ versus $H_1: \theta \in \Theta_0^c$ is given by

$$\lambda_{LR} = -2 \log \frac{\sup_{\theta \in \Theta_0} L(\theta)}{\sup_{\theta \in \Theta} L(\theta)},$$

where $\theta$ is a population parameter, $\Theta$ is the entire parameter space, $\Theta_0$ is a subset of $\Theta$, and $\Theta_0^c$ is the complement of $\Theta_0$. If maximum likelihood estimates $\hat{\theta}$ and $\hat{\theta}_0$ exist, we have

$$\lambda_{LR} = -2 \log \frac{L(\hat{\theta}_0)}{L(\hat{\theta})}.$$

By Wilks' theorem, under the null model, the test statistic $\lambda_{LR}$ is asymptotically $\chi^2$-distributed with degrees of freedom corresponding to the difference in dimensionality of $\Theta$ and $\Theta_0$ [13]. The number of degrees of freedom is one for all three modeling approaches. As in the previous interaction molecular QTL mapping studies [14, 7], we performed multiple testing correction using the Benjamini–Hochberg method [15] with a false discovery rate (FDR) threshold of 0.1.

#### 2.5 Evaluation metrics for simulation experiments

Model performance was evaluated using four metrics:

- Calibration of $P$ values: using synthetic data obtained from the interaction null model, which contains the genotype and treatment terms but lacks the G×T interaction term.
- Statistical power: using synthetic data obtained from the full model including the G×T interaction term with varying effect sizes.

- Overall performance: we computed the true positive rate (TPR) as well as the false discovery rate (FDR) using the synthetic data consisting of realistic proportions of data-generating models.

- Quality of effect estimation: we compared the estimated effects against the true values used for data generation.

#### 2.6 Generating data for simulations

For simulation experiments, we generated $10,000$ feature-SNP pairs from the nonlinear regression model in Eq. (2) for each analysis, using the `make_sim_data()` function. Specifically, we generated paired and unpaired data for assessing calibration of $P$ values, power to detect non-zero G×T interactions, and overall performance based on true and false positive rates. In the paired data, each simulated feature-SNP pair comprised the genotypes for 78 individuals and simulated transformed count data $\mathbf{y}^* = \{y_i^*\}_{i=1}^{156}$ (i.e., the phenotype vector), corresponding to 78 phenotype values for each of the control and treated conditions (156 in total). In the unpaired data, each simulated feature-SNP pair comprised the genotypes for 156 individuals and simulated transformed count data $\mathbf{y}^* = \{y_i^*\}_{i=1}^{156}$. Among the individuals, 93 and 63 were in control and treated conditions, respectively. The genotypes were drawn from a Binomial distribution, $\text{Binom}(n = 2, p = \pi)$, where $\pi \sim \text{Uniform}(0.05, 0.5)$, corresponding to minor allele frequency (MAF) ranging from 0.05 to 0.5. Thus, $g = 0$ and $g = 2$ correspond to the major and minor allele homozygotes, respectively. In generating a simulated phenotype vector from Eq. (2), we fixed the intercept and residual error standard deviation to $\beta_0 = 0$ and $\sigma = 1$. The regression coefficients for the genotype and treatment terms were drawn from Normal distributions as $\beta_g \mid \sigma^2 \sim \mathcal{N}(0, \phi_g^2\sigma^2)$ and $\beta_t \mid \sigma^2 \sim \mathcal{N}(0, \phi_t^2\sigma^2)$ with $\phi_g = 1.5$ and $\phi_t = 2.0$, respectively, unless otherwise noted.

##### 2.6.1 Simulations focusing on AA-AB-BB SNPs

For the analysis focusing on AA-AB-BB SNPs (i.e., SNPs with all three genotypes observed), we excluded the AA-AB SNPs (i.e., SNPs missing the minor allele homozygote) by setting the input argument `filter.geno=TRUE` when calling the `make_sim_data()` function. For assessing the calibration, we generated $10,000$ feature-SNP pairs, setting $\beta_{g\times t} = 0$. For evaluating the statistical power, we fixed the interaction effect to $\beta_{g\times t} \in \{0.25, 0.50, 0.75, 1.00\}$ for each set of $10,000$ feature-SNP pairs. These values were based on the results of model fitting and parameter estimation using the hNPC eQTL data [7]. For evaluating the overall performance by computing true positive rate (TPR) and false discovery rate (FDR), we generated 200 feature-SNP pairs, setting $\beta_{g\times t} = 1.0$. For the remaining $9,800$ pairs, we set $\beta_{g\times t} = 0$ (10,000 in total).

##### 2.6.2 Simulations focusing on AA-AB SNPs

For the analysis of the AA-AB SNPs (Sections 3.3–3.5), we included the AA-AB SNPs by setting `filter.geno=FALSE` when calling the `make_sim_data()` function. The 10,000 simulated data contained 1,223 and 1,773 AA-AB SNPs for the paired and unpaired settings, respectively. For illustrating the optimization problem for the AA-AB SNPs (Sections 3.3 and 3.4), we fixed $\beta_g$, $\beta_t$, and $\beta_{g\times t}$ at values summarized in Table S2. For assessing general parameter estimation quality, we drew the G×T interaction effect as $\beta_{g\times t} \mid \sigma^2 \sim \mathcal{N}(0, \phi_{g\times t}^2\sigma^2)$ where $\phi_{g\times t} = 1.0$.

##### 2.6.3 Kinship matrices for unpaired data from the 1000 Genomes Project

For the unpaired data, we constructed genomic relationship matrices (GRMs) from 312 randomly selected samples in African (AFR), European (EUR), East Asian (EAS), and South Asian (SAS) ancestry populations from the 1000 Genomes Project [16], including second-degree relatives. We used the upper-left 156 by 156 and 312 by 312 matrices as kinship matrices for data generation and model fitting. We evaluated the models using each of the four GRMs as a kinship matrix. For other analyses, we focused on the GRM from the AFR population.

#### 2.7 Software

All statistical and computational analyses were conducted with the R statistical programming language [10] and the Snakemake workflow [17]. Portions of code were generated with the assistance of Claude Code (Anthropic) and Codex (OpenAI). We also used Claude (Anthropic) and ChatGPT-5 (OpenAI) to improve documentation quality. We reviewed and verified the materials produced by these tools.

### 3 Supplementary Results

#### 3.1 Simulations for AA-AB-BB SNPs

In simulation experiments, we considered six scenarios by generating data with three levels of random effect standard deviation (zero, medium, and high) and fitting models with or without a random effect term (see Table S1). Specifically, the six scenarios were:

1. no random effect,
2. an intermediate-level random effect in data generation but no random effect term in model fitting,
3. a high-level random effect in data generation but no random effect term in model fitting,
4. no random effect in data generation but with a random effect term in model fitting,
5. an intermediate-level random effect in data generation and a random effect term in model fitting,
6. a high-level random effect in data generation and a random effect term in model fitting.

We generated the data based on the nonlinear model and compared the three modeling approaches, nonlinear, linear, and RINT, based on $P$ value calibration, power, and FDR-TPR plots. We performed analyses in the paired setting, where the data contain one sample per combination of genotype and condition, and in the unpaired setting, where the data contain one sample from each donor.

##### 3.1.1 Paired setting

For the paired setting, $P$ value calibration, power, and FDR-TPR plots are shown in Figs. S17, S18, and S19, respectively.

##### 3.1.2 Unpaired setting

For the unpaired setting, we repeated the analysis using GRMs constructed from samples in four ancestry populations, AFR, EUR, EAS, and SAS. $P$ value calibration plots are shown in Figs. S26, S27, S28, and S29. Power plots are shown in Figs. S30, S31, S32, and S33. FDR-TPR plots are shown in Figs. S34, S35, S36, and S37.

#### 3.2 Application to eQTL mapping data in hNPCs

Fig. S44 compares interaction eQTLs in hNPCs with and without growth stimulation obtained by a previous study and those obtained by the nonlinear, linear, and RINT approaches.

#### 3.3 On the practical non-identifiability for certain AA-AB SNPs

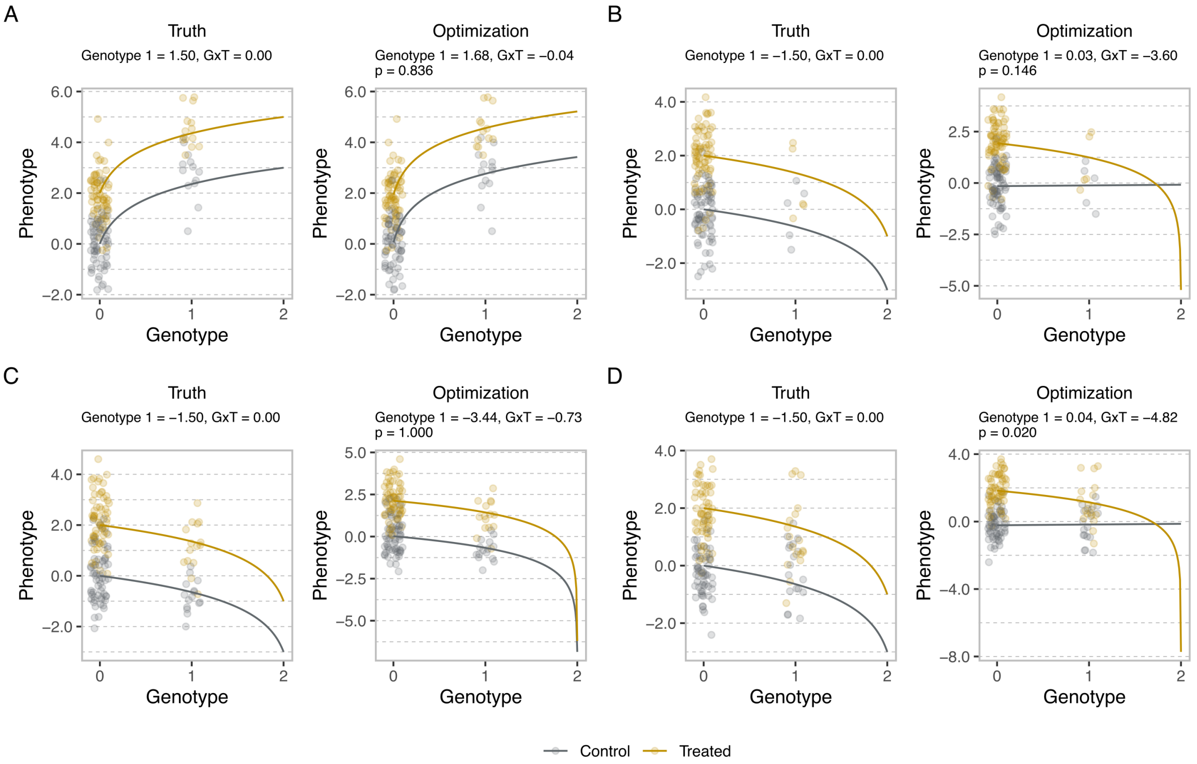

Figure S1: An example of simulated molecular count data in control and treated conditions in case 1 (A) and case 2 (B–D). The left panel shows functional relationships between genotypes and phenotypes based on the true values used in data generation, whereas the right panels show fitted curves from nonlinear regression.

In this section, we illustrate the problem of estimating genotype effects by the nonlinear approach for some of the AA-AB SNPs using simulated data. The key idea is that estimation of the genotype effect is hampered when data points are present only in the flat portion of the nonlinear regression curve, as we explain below. For concreteness, we generated data based on the nonlinear model, fitted the nonlinear models, and computed LRT $P$ values. For simplicity, we omitted random effects and focused on paired data. We considered four representative data generation cases with the genotype, treatment, and G×T interaction effects fixed at different values, as summarized in Table S2. In cases 1 and 2, the interaction effect was set to zero, whereas it was set to non-zero in cases 3 and 4. In cases 1 and 3, the genotype effects in both control and treated conditions were set to positive values, whereas they were set to negative values in cases 2 and 4. For each case, we drew a MAF from a uniform distribution on $[0.05, 0.5]$ and then genotypes from a binomial distribution with size two and probability equal to the MAF to obtain multiple examples of the AA-AB SNPs for generating phenotypes. This ensured that $g = 0$ and $g = 2$ were assigned to the major and minor allele homozygotes, as in other simulations in this work.

Fig. S1 shows illustrative examples of the AA-AB SNPs in case 1 (A) and case 2 (B–D). Here, the $x$-axis values represent genotypes $g$, and the differences in the $y$-axis values between $g = 0$ and $g = 2$ correspond to the genotype effect $2\beta_g$ and $2\beta_g + 2\beta_{g\times t}$ for the control and treated conditions, respectively. The lines in the left and right panels are based on true parameters and nonlinear regression estimates, respectively, overlaid on the same data.

##### 3.3.1 AA-AB SNPs when G×T is absent

**Reasonable identifiability when data cover the steep/lower part of the phenotype-genotype curve.** Fig. S1A shows an example in case 1, where the true genotype effects were positive. Notably, data points were present in the steep portion of the nonlinear regression curve, as these correspond to major allele homozygotes and heterozygotes. The output values from the `map_qtl()` function with nonlinear regression were 1.68 and −0.04 for the genotype effect in the control condition and the G×T interaction effect, respectively, which correspond to the genotype effect of 1.64 in the treated condition. The nominal $P$ value from LRT was 0.836, consistent with the data being generated with a zero interaction effect. Overall, these represent reasonable optimization results.

**Poor identifiability when data are only in the flat/upper part of the phenotype-genotype curve.** By contrast, Fig. S1B–D show examples in case 2, where the true genotype effects were negative. Notably, data points were present only in the flat portion of the nonlinear regression curve. In these cases, the output values from the `map_qtl()` function with nonlinear regression for the genotype effect in the control condition and the G×T interaction effect were largely discrepant from the true values. We note that the numbers returned by the function as nominal $P$ values from LRT (0.145, 0.999, and 0.020) may not be valid, as the genotype effect estimates from optimization may not represent the MLE.

##### 3.3.2 AA-AB SNPs when G×T is present

Fig. S2 shows four illustrative examples of the AA-AB SNPs in case 3 (A) and case 4 (B–D), where the true G×T interaction effect was non-zero.

**Reasonable identifiability when data cover the steep/lower part of the phenotype-genotype curve.** Fig. S2A shows an example in case 3, where the true genotype effects were positive. As in case 1, data points were present in the steep portion of the nonlinear regression

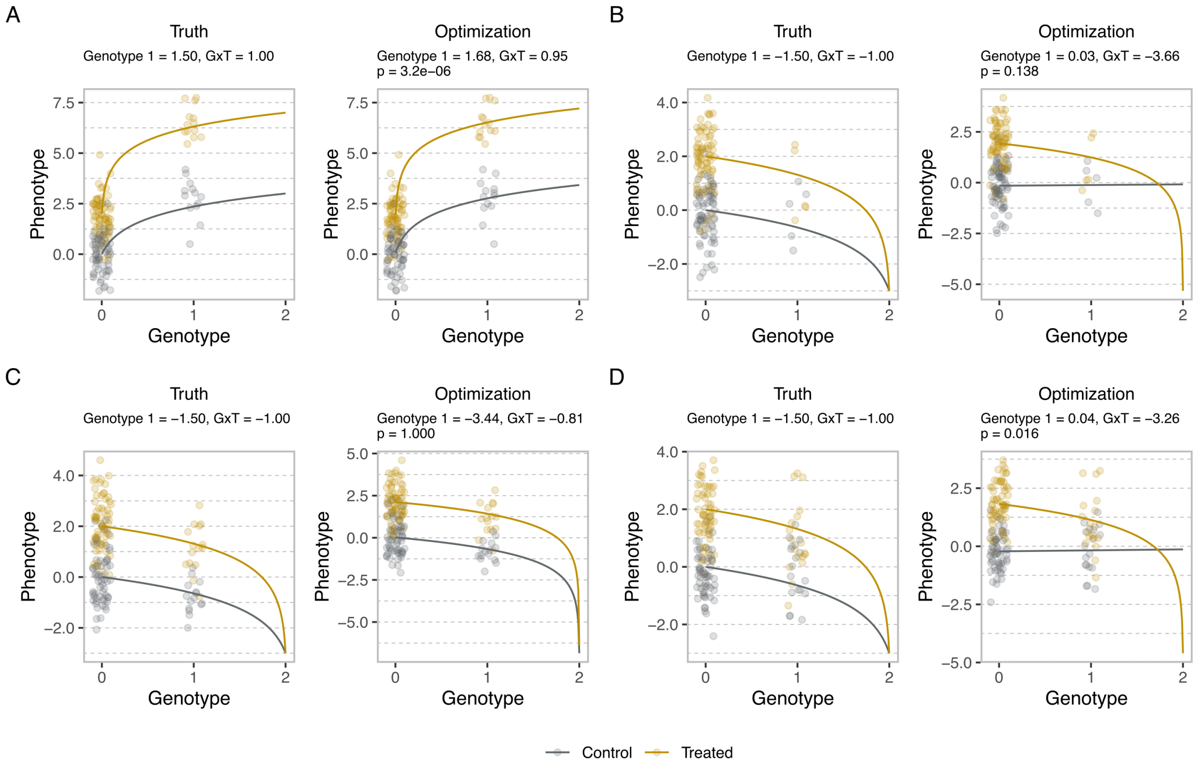

Figure S2: An example of simulated molecular count data in control and treated conditions in case 3 (A) and case 4 (B–D). The left panel shows functional relationships between genotypes and phenotypes based on the true values used in data generation, whereas the right panels show fitted curves from nonlinear regression.

curve, as these correspond to major allele homozygotes and heterozygotes. The output values from the `map_qtl()` function with nonlinear regression were 1.68 and 0.95 for the genotype effect in the control condition and the G×T interaction effect, respectively, which correspond to the genotype effect of 2.63 in the treated condition. The nominal $P$ value from LRT was $3.2 \times 10^{-6}$, consistent with the data being generated with a non-zero interaction effect. Overall, these represent reasonable optimization results.

**Poor identifiability when data are only in the flat/upper part of the phenotype-genotype curve.** By contrast, Fig. S2B–D show examples in case 4, where the true genotype effects were negative. As in case 2, data points were present only in the flat portion of the nonlinear regression curve. In these cases, the output values from the `map_qtl()` function with nonlinear regression for the genotype effect in the control condition and the G×T interaction effect were largely discrepant from the true values. As in case 2, the numbers returned by the function as nominal $P$ values from LRT (0.138, 1.000, and 0.016) may not be valid, as the genotype effect estimates from optimization may not represent the MLE.

##### 3.3.3 Practical non-identifiability and degradation of estimates

The above results are consistent with the idea that, with nonlinear regression, genotype effects are "practically non-identifiable" when data are absent from the steep portion of the curve. This is likely due to a wide range of parameter values giving similar likelihood, as the data do not contain sufficient information about the curvature. This can also lead an optimization algorithm to terminate at values far from the MLE. This type of "degradation of estimates" has also been reported previously for SNPs with low minor allele frequency [1]. We note that linear regression does not encounter the issue of practical non-identifiability. However, the genotype effect estimates were more strongly biased for the AA-AB SNPs where true values were near zero or negative than for other SNPs (see the third column in Figs. S3 and S4). The RINT approach performed poorly across the cases with estimates that largely differed from the true values (see the fourth column in Figs. S3 and S4).

#### 3.4 Calibration and power for the AA-AB SNPs

To systematically assess calibration and power for the AA-AB SNPs, we generated and fitted the models for 10,000 feature-SNP pairs in all four cases for both paired and unpaired data. We first assessed the parameter estimation quality by visually comparing the optimization output and the true parameter values separately for the AA-AB-BB and AA-AB SNPs for paired (Figs. S5–S8) and unpaired data (Figs. S9–S12). In cases 1 and 3, nonlinear regression yielded approximately unbiased genotype effect estimates for both AA-AB-BB and AA-AB SNPs. In cases 2 and 4, nonlinear regression yielded genotype effect estimates that were largely discrepant from the true values for AA-AB SNPs, while estimates remained approximately unbiased for AA-AB-BB SNPs. Unlike the nonlinear approach, the linear and RINT approaches produced systematically biased genotype effect estimates for both AA-AB-BB and AA-AB SNPs (Figs. S5–S12).

The issue with nonlinear regression for AA-AB SNPs in cases 2 and 4 was also evident from the poorly calibrated $P$ values, which were skewed toward one (Figs. S13D and S14D), and the low power (Figs. S13E and S14E) for both paired and unpaired data. This is expected as a substantial fraction of $P$ values are unlikely to be valid, as we discussed in the previous section (Section 3.3). While the linear approach yielded well-calibrated $P$ values in cases 1 and 2 for both AA-AB-BB and AA-AB SNPs when the data were paired (Fig. S13A–D), the power to detect significant G×T interactions for AA-AB SNPs was low in case 4 (Fig. S13E). For unpaired data, the linear approach resulted in anti-conservative behavior (Fig. S14A–D). The RINT approach performed poorly across cases and data types.

#### 3.5 General assessment of parameter estimation quality

To illustrate the practical non-identifiability of the nonlinear regression model for certain AA-AB SNPs more generally, we generated and fitted the models for 10,000 feature-SNP pairs in the six scenarios for each of the paired and unpaired settings with varying rather than fixed parameter values (Section 2.6) and visually compared the optimization output and the true values separately for the AA-AB-BB and AA-AB SNPs (Figs. S20–S25 and S38–S43). As we note in the main text, for the AA-AB-BB SNPs, the nonlinear approach yielded substantially less biased estimates of the genotype effects in the control and treated conditions ($\beta_g$ and $\beta_g + \beta_{g\times t}$) than the linear and RINT approaches (Figs. S20–S25A and S38–S43A). By contrast, for some of the AA-AB SNPs, the optimization algorithm returned unrealistic values for the genotype effects with strong deviation from the true genotype effects (Figs. S20–S25B and S38–S43B). Notably, the unrealistic values were returned when the true genotype effects were near zero or negative. As we assigned the major and minor alleles to $g = 0$ and $g = 2$, these correspond to situations where data points are present only in the flat portion of the curve ($g = 0$ and $g = 1$) as we illustrated earlier.

##### 3.6 Frequency of the AA-AB SNPs

In this section, we elaborate on the frequency of AA-AB SNPs in the eQTL data in hNPCs [7]. As we described in the main text, among 3,401 gene-SNP pairs on autosomes in the union of condition-stratified eQTL mapping, 3,083 pairs contained the AA-AB-BB SNPs. To corroborate this observation, assuming HWE, we calculated the probability that a genetic variant is an AA-AB SNP based on

$$\Pr(\text{AA-AB}) = (1 - \pi^2)^m,$$

where $\pi$ and $m$ denote minor allele frequency and the number of subjects, respectively. Table S3 shows the probability Pr(AA-AB) at representative MAFs, ranging from 0.01 to 0.5, when the number of subjects was 78 as in the hNPC data (see Section 2.1). The MAF ranges are relevant, as SNPs with MAF ranging from 0.01 to 0.5 without significant departure from Hardy–Weinberg equilibrium (HWE) were selected prior to condition-stratified eQTL mapping in hNPCs. Notably, Pr(AA-AB) equaled 0.992 and 0.457 at MAF of 0.01 and 0.1, respectively, while the probability was only 0.041 at MAF of 0.2. However, in the union set of condition-stratified eQTLs in hNPCs, corresponding to 3,220 unique SNPs, the number of SNPs with MAF ranging from 0.01 to 0.1 was smaller than that in other MAF ranges, likely due to challenges in detecting associations at low MAF (Table S4), resulting in the relatively low overall frequency of the AA-AB SNPs. Similar situations are expected for molecular QTL data in general, as the eQTL analysis in hNPCs follows the standard procedure.

##### 3.7 Computational performance

To systematically evaluate the computational performance of our method, we measured the runtime and memory usage of the `map_qtl()` function on a Linux-based system. In this series of analyses, we simulated $10,000$ paired and unpaired data in the same manner as for the computation of TPR and FDR. We considered all six scenarios and repeated measurements five times. In all instances, the memory usage was less than 0.5 gigabytes (Table S5). For the paired data, the maximum runtime across scenarios, modeling approaches, and replicates was less than 2,000 and 5,600 seconds at the sample sizes of 156 and 312, respectively (Fig. S15). For the unpaired data, the maximum runtime was less than 2,300 and 5,600 seconds at the sample sizes of 156 and 312, respectively (Fig. S16). These results suggest that our software provides adequate computational efficiency.

#### 4 Supplementary Discussion

We developed a software package to facilitate molecular interaction QTL mapping accounting for allelic additivity and correlated samples. Using this package, we examined the impacts of these modeling choices on simulated data. Specifically, we used simulations to compare nonlinear regression based on the allelic additivity assumption with two modeling approaches, log transformation followed by linear regression with an additive genotype variable and RINT transformation followed by linear regression. We also examined the effects of accounting for correlation structure by including donor or polygenic random effects. We considered paired data, which contained repeated measurements for a given donor, and unpaired data, which contained exactly one sample per donor. Our analyses suggest that the nonlinear approach outperforms the linear and RINT approaches for both paired and unpaired data and that modeling donor random effects is crucial for maintaining the power to detect G×T interactions. In our reanalysis of previously published hNPC eQTL data, we obtained overlapping but distinct sets of SNPs with significant G×T interactions using the nonlinear, linear, and RINT approaches. Although the ground truth is unknown, given our previous observation that nonlinear regression captures the data better than linear regression [6], we interpreted the differences in the sets as indicating improved accuracy of the nonlinear approach. Overall, the results suggest that it is beneficial to use nonlinear regression rather than linear regression to model the relationship between genotypes and molecular phenotypes in interaction molecular QTL mapping.

A limitation of our nonlinear approach is that the AA-AB SNPs need to be excluded. As we illustrated in Section 3.3, in case 4, where data were absent from the steep portion of the curve, the genotype effects were practically non-identifiable, often resulting in instability in the optimization algorithm (Figs. S5–S12). Although the linear and RINT approaches did not result in instability, they were ineffective in detecting G×T interactions in simulated data with non-zero effect under the allelic additivity assumption (Figs. S13E and S14E). We argue that excluding this group of SNPs is well justified. By contrast, it is not ideal to discard the AA-AB SNPs in case 3, for which data were present in the steep portion of the curve. However, unlike in simulated data, it is not straightforward to distinguish between situations similar to case 3 and those similar to case 4. Thus, it can be justified to exclude all AA-AB SNPs, as the overall fraction of these SNPs is expected to be relatively small in typical molecular QTL mapping datasets, which focus on common alleles. Nevertheless, in principle, AA-AB SNPs in a case 3-like situation can be recovered by assessing the practical non-identifiability of the genotype effects based on the widths of the finite sample profile likelihood confidence intervals [18]. Alternatively, the linear approach can be applied to the AA-AB SNPs.

There are multiple future directions to consider. One particular concern is whether to include both donor and polygenic random effects for paired data, as opposed to including only one of them in the models. DetectGxT currently allows zero or one random effect term for computational simplicity and efficiency. We propose to include donor random effects, as polygenicity and population stratification can be accounted for by residualizing the phenotype against genotype PCs prior to model fitting. However, if the samples contain known or cryptic relatedness, it is desirable to account for it by including a polygenic random effect term to avoid spurious associations. In principle, it is possible to include both donor and polygenic random effects, which can potentially perform better than our current approach. Indeed, a previous study has developed an interaction molecular QTL mapping approach, termed “suez,” that accounts for both donor and polygenic random effects, as well as latent confounding factors [19]. In the suez method, a global covariance matrix that includes subject and polygenic random effects as well as latent confounding is estimated via optimization across features prior to fitting models for individual feature-SNP pairs. It can be useful to incorporate the nonlinear regression model into the suez method.

Another future direction is to systematically compare DetectGxT to approaches using generalized linear models (GLMs), which fit molecular count data on the original scale, in the context of G×T or gene-by-environment (G×E) interactions as well as in bulk and pseudobulk single-cell data.

Although we chose to model the data with homoscedastic Gaussian noise on the transformed scale for computational efficiency, recent studies found effectiveness of negative binomial GLMs for bulk and pseudobulk molecular QTL mapping [20, 21]. Notably, another study performed continuous state-dependent eQTL mapping in single-cell data employing Poisson mixed-effects models [22]. However, we also note that these studies used a linear predictor to model the relationship between genotypes and phenotypes without accounting for the potentially nontrivial model misspecification due to the log link, which may have resulted in inaccurate inference. Therefore, to allow for formal comparison between modeling original count and transformed data, it would be of interest to develop a generalized linear regression model where the linear predictor is replaced with our nonlinear function. Indeed, for a single-condition eQTL mapping, a pioneering study has developed the TReC model, which uses a modified negative binomial model where the linear predictor is replaced with a weighted function to account for the nonlinear relationship [23].

Finally, we note that, although we focused on the two-step procedure here, Knowles et al. and Kasela et al. have successfully identified a large number of interaction eQTLs in an in vitro cell system and tissue samples, respectively, using a one-step procedure where all *cis*-SNPs proximal to genes were tested for G×E interaction as in single-condition (or condition-stratified) molecular QTL mapping [19, 24]. It would be of interest to apply this method to the hNPC and other existing datasets and systematically compare the results with those from the two-step procedure. Although the two-step procedure is commonly used in G×E analysis for general continuous phenotypes to facilitate the detection of interactions [25, 26], the one-step procedure can be generally effective for G×E or G×T analysis on molecular count phenotypes where effect sizes are typically larger. In the one-step procedure, as in the study by Knowles et al. [19], SNP filtering and multiple testing schemes established for single-condition molecular QTL mapping can readily be applied [27], which can further improve the accuracy of G×T analysis on molecular count phenotypes.

### 5 Supplementary Tables

Table S1: Simulation scenarios.

| Scenario | Data generation | Model fitting |
|---|---|---|
| 1 | Zero | Excluded |
| 2 | Medium | Excluded |
| 3 | High | Excluded |
| 4 | Zero | Included |
| 5 | Medium | Included |
| 6 | High | Included |

The **Data generation** column indicates the level of random effect standard deviation in simulating data. The **Model fitting** column indicates whether the random effect term was included when fitting the data.

Table S2: Cases for illustration.

| Case | Genotype 1 | G×T | Genotype 2 |
|---|---|---|---|
| 1 | 1.5 | 0.0 | 1.5 |
| 2 | -1.5 | 0.0 | -1.5 |
| 3 | 1.5 | 1.0 | 2.5 |
| 4 | -1.5 | -1.0 | -2.5 |

The **Genotype 1** and **G×T** columns contain the genotype effect in the control condition $\beta_g$ and G×T interaction effect $\beta_{g\times t}$, respectively. The **Genotype 2** column contains the genotype effect in the treatment condition, which corresponds to $\beta_g + \beta_{g\times t}$. Note that the intercept $\beta_0$, treatment effect $\beta_t$, and residual error standard deviation $\sigma$ were fixed at 0.0, 2.0, and 1.0.

Table S3: Probability that a genetic variant is an AA-AB SNP.

| MAF | Pr(AA-AB) |
|---|---|
| 0.01 | 0.992 |
| 0.05 | 0.823 |
| 0.10 | 0.457 |
| 0.20 | 0.041 |
| 0.30 | $6.39 \times 10^{-4}$ |
| 0.40 | $1.24 \times 10^{-6}$ |
| 0.50 | $1.80 \times 10^{-10}$ |

Shown is the theoretical probability that a genetic variant is an AA-AB SNP at a given MAF. The number of subjects was set to 78.

Table S4: Distribution of MAF in the hNPC eQTL data.

| MAF range | Number of SNPs |
|---|---|
| [0.01, 0.1] | 346 |
| (0.1, 0.2] | 640 |
| (0.2, 0.3] | 694 |
| (0.3, 0.4] | 769 |
| (0.4, 0.5] | 771 |

Shown are the numbers of unique SNPs in MAF bins in the hNPC eQTL data. The total number of unique SNPs was 3,220.

Table S5: Runtime and memory usage.

| Data | Sample size | Method | Runtime | Memory |
|---|---|---|---|---|
| Paired | 156 | Nonlinear | 1970.600 | 425.270 |
| Paired | 156 | Linear | 726.186 | 427.800 |
| Paired | 156 | RINT | 780.301 | 431.410 |
| Paired | 312 | Nonlinear | 5503.699 | 451.230 |
| Paired | 312 | Linear | 1810.960 | 439.970 |
| Paired | 312 | RINT | 2034.756 | 446.070 |
| Unpaired | 156 | Nonlinear | 1974.452 | 427.750 |
| Unpaired | 156 | Linear | 1483.463 | 435.350 |
| Unpaired | 156 | RINT | 2220.002 | 440.040 |
| Unpaired | 312 | Nonlinear | 5513.914 | 440.450 |
| Unpaired | 312 | Linear | 4486.450 | 454.260 |
| Unpaired | 312 | RINT | 4370.025 | 449.880 |

Shown are the runtime and memory usage of the `map_qtl()` function for 10,000 feature-SNP pairs. The **Data** column indicates the type of data. The **Method** column indicates modeling approaches used in the analysis. The **Runtime** column contains the maximum values of the runtime in seconds across five replicates and six scenarios. The **Memory** column contains the maximum values of the maximum resident set size in megabytes.

### 6 Supplementary Figures

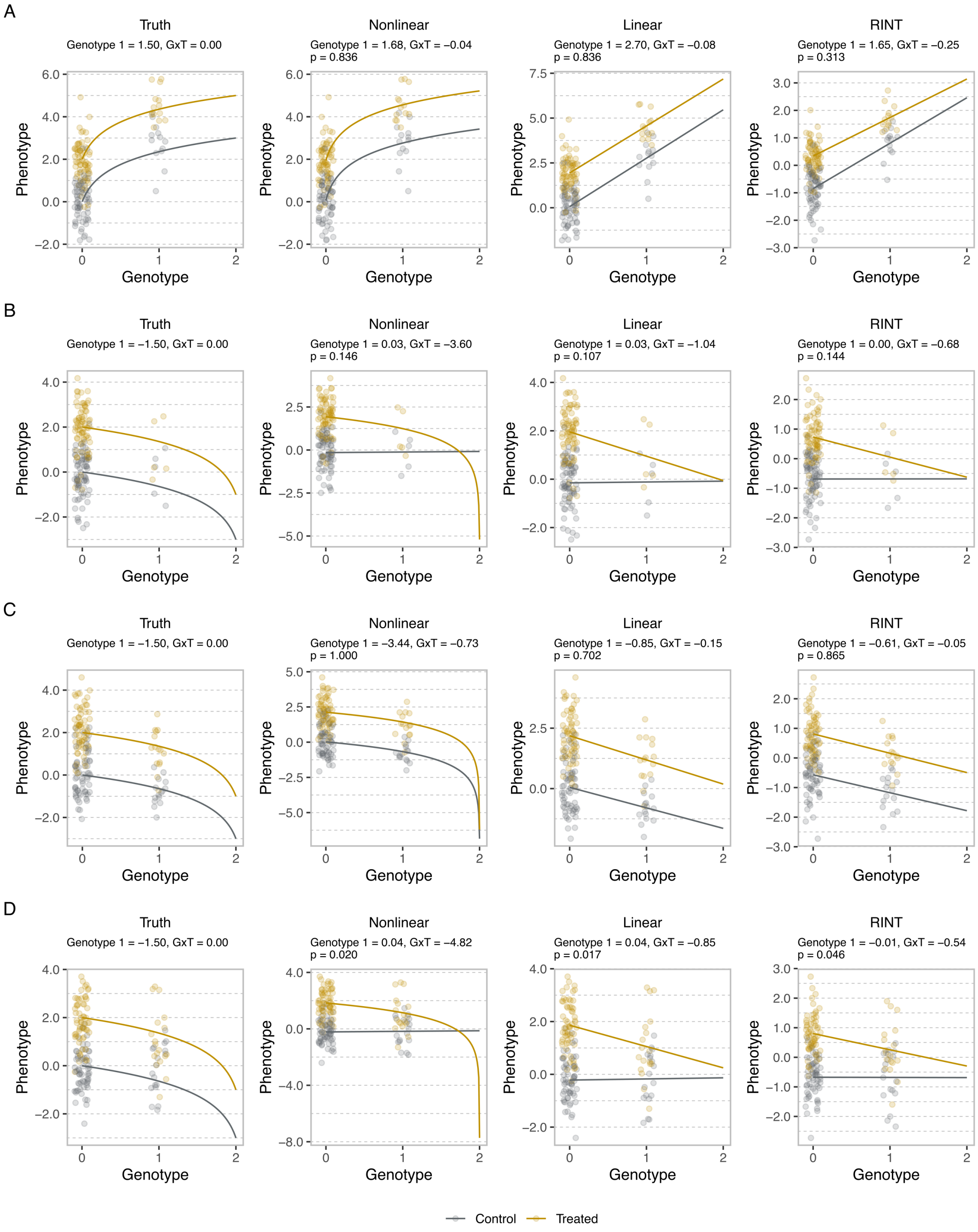

Figure S3: An example of simulated molecular count data in control and treated conditions in case 1 (A) and case 2 (B–D). The first column shows functional relationships between genotypes and phenotypes based on the true values used in data generation, whereas the second to fourth columns show fitted curves from the nonlinear, linear, and RINT approaches, respectively.

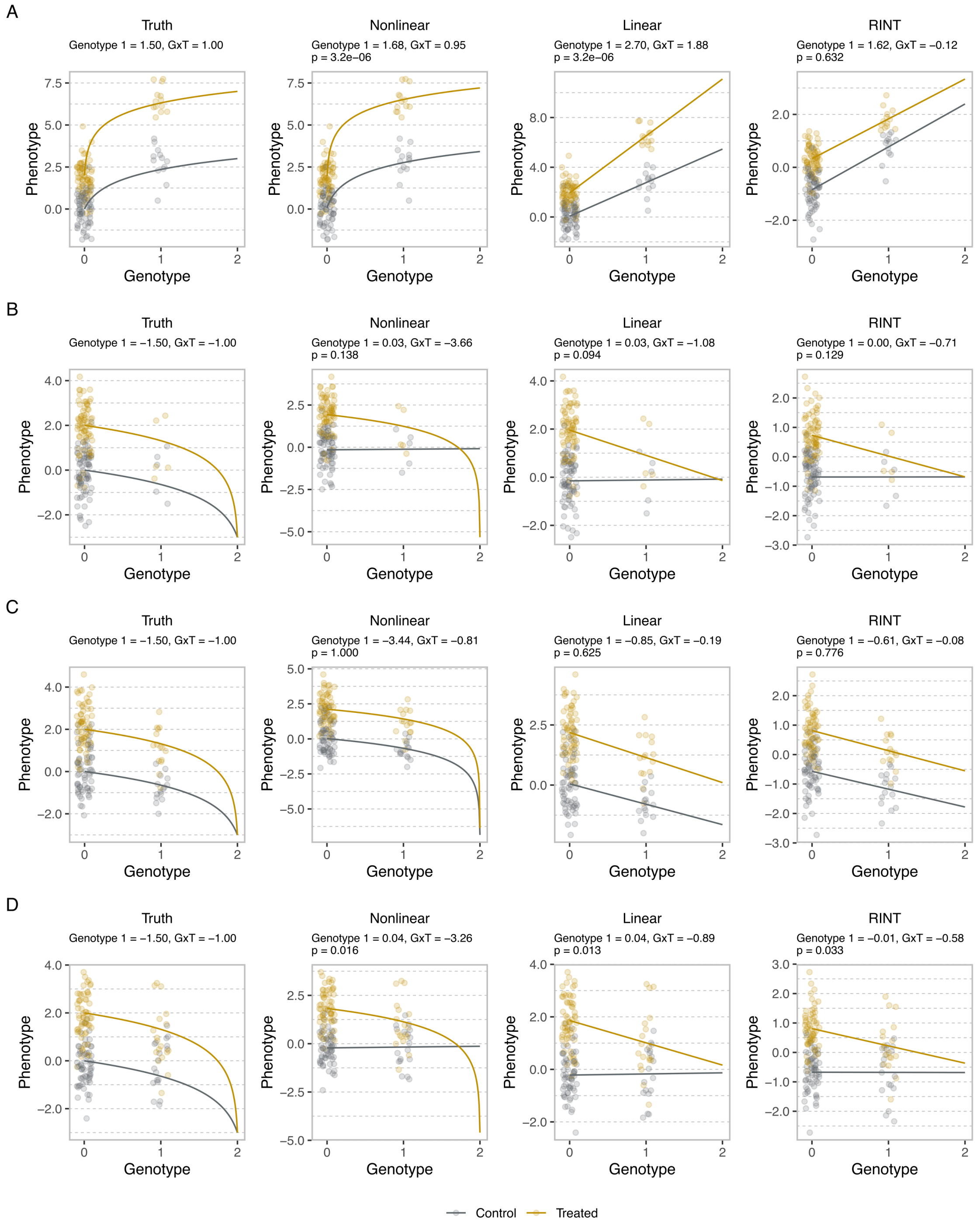

Figure S4: The same as Fig. S3 but for case 3 (A) and case 4 (B–D).

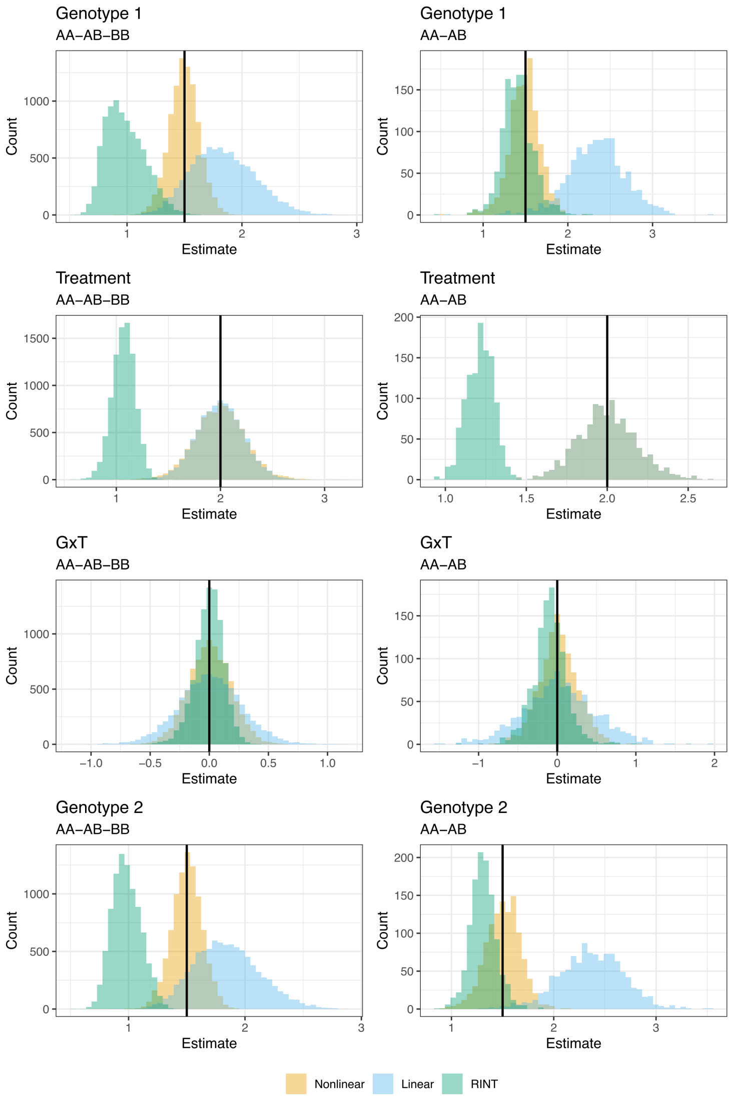

Figure S5: Histograms of effect estimates in case 1. The vertical lines represent true values. The labels Genotype 1, Treatment, G×T, and Genotype 2 represent genotype effects in the control condition, treatment effects, G×T interaction effects, and genotype effects in the treated condition, respectively. The orange, blue, and green colors represent the nonlinear, linear, and RINT approaches, respectively. The left and right panels show results for the AA-AB-BB and AA-AB SNPs, respectively.

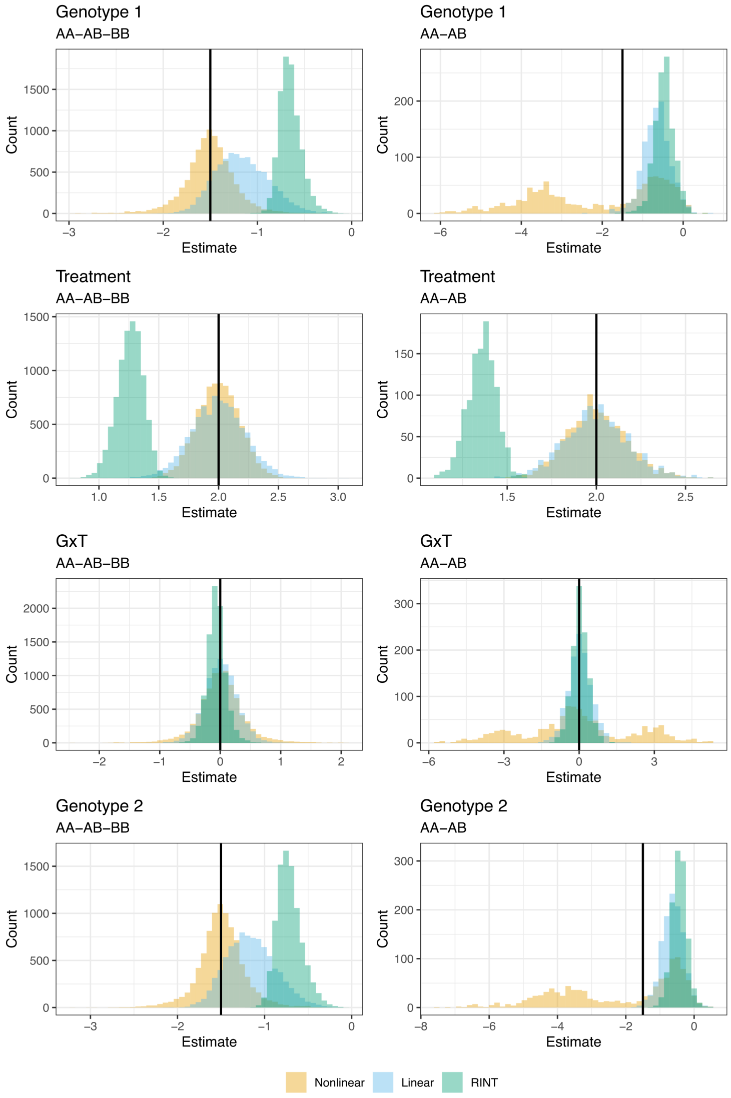

Figure S6: The same as Fig. S5 but in case 2.

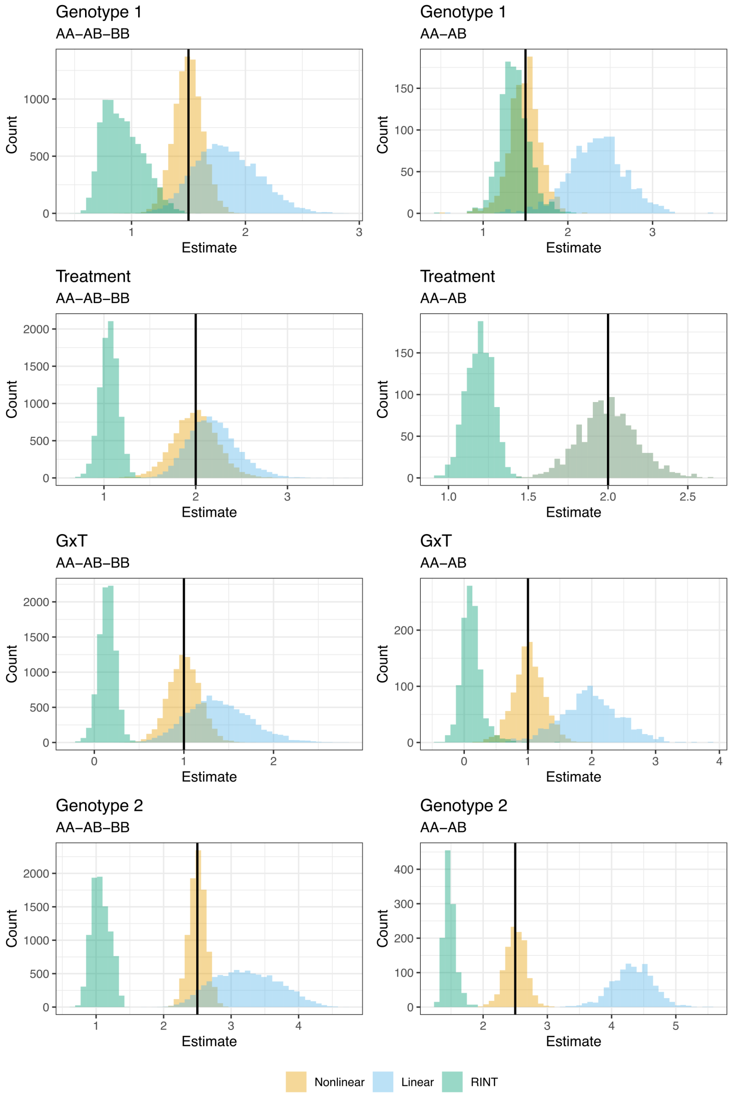

Figure S7: The same as Fig. S5 but in case 3.

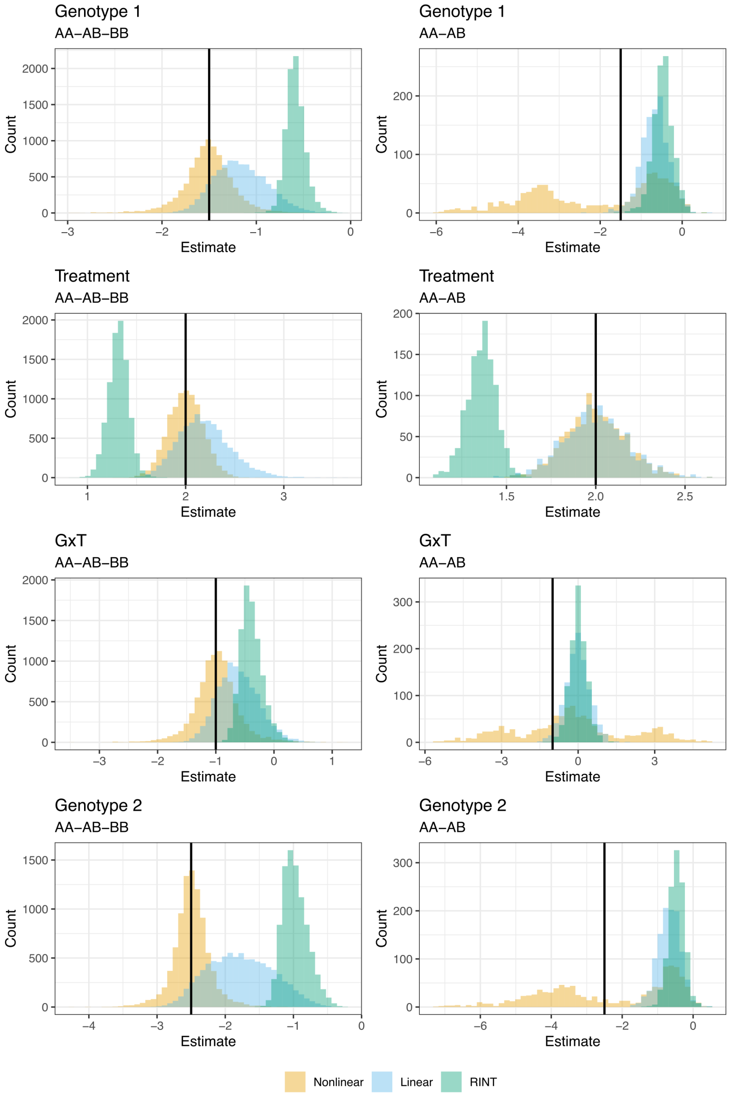

Figure S8: The same as Fig. S5 but in case 4.

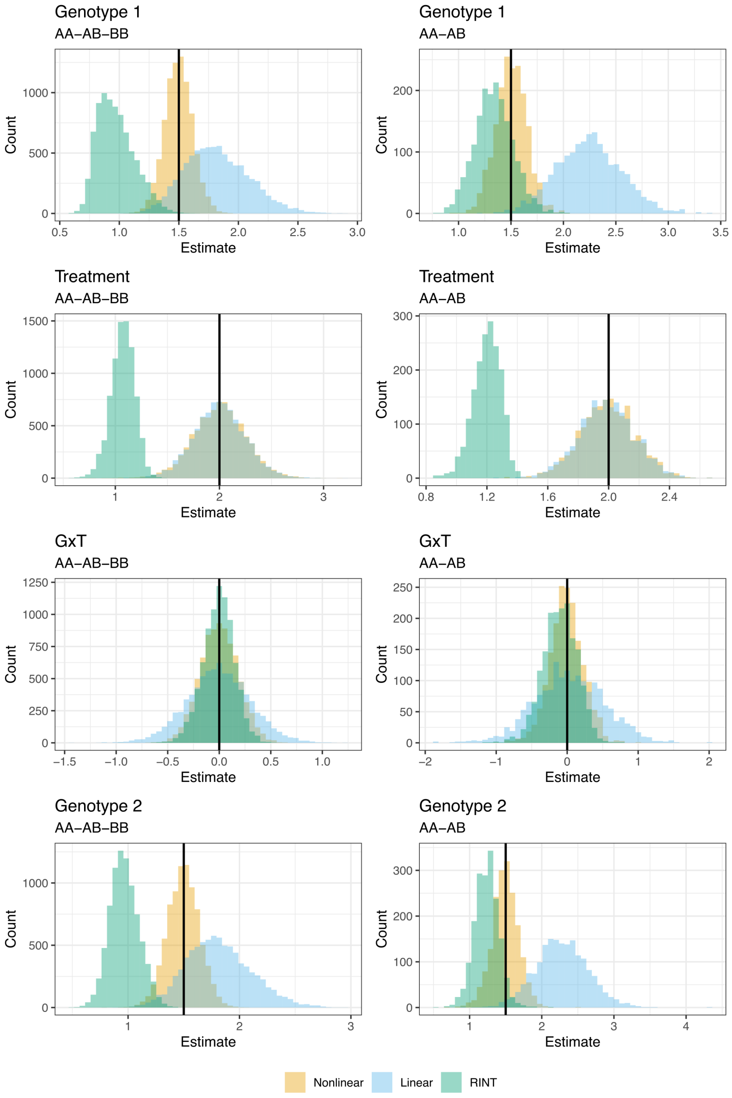

Figure S9: The same as Fig. S5 but for unpaired data.

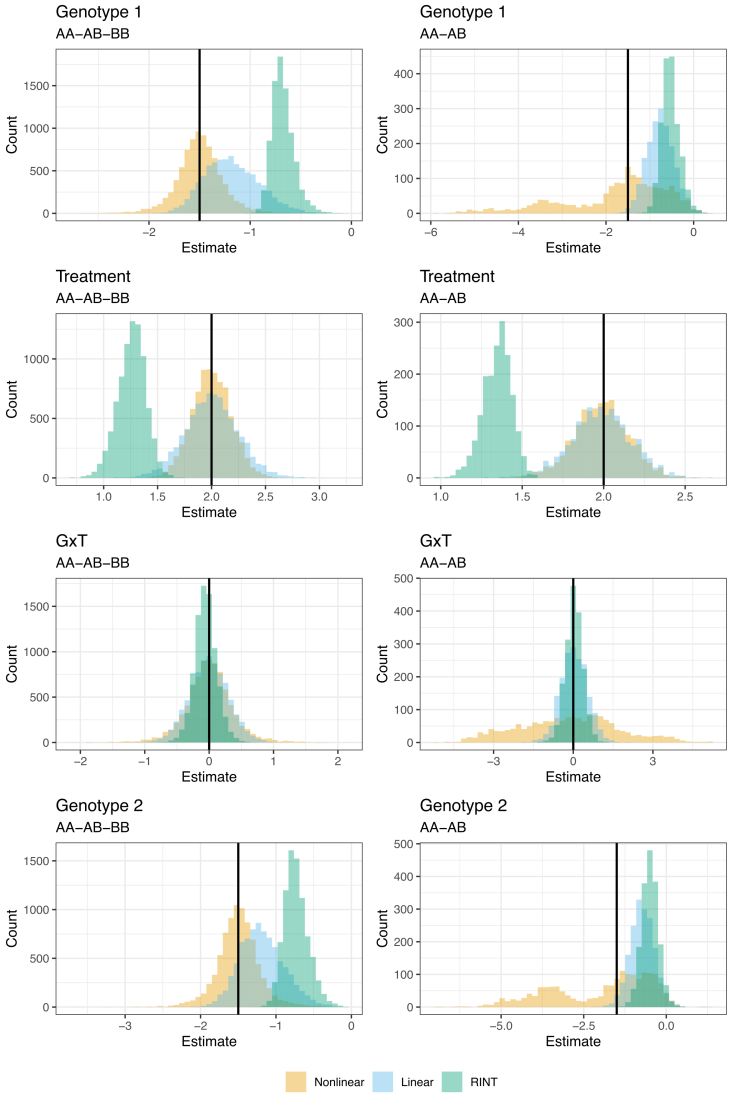

Figure S10: The same as Fig. S6 but for unpaired data.

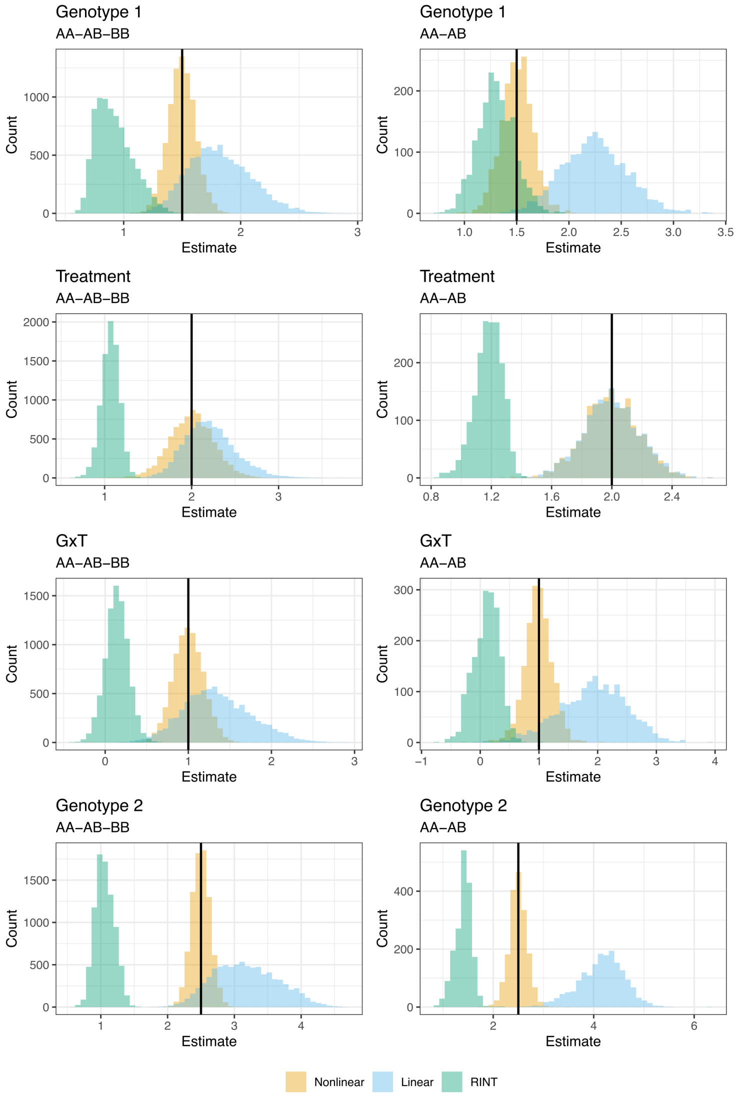

Figure S11: The same as Fig. S7 but for unpaired data.

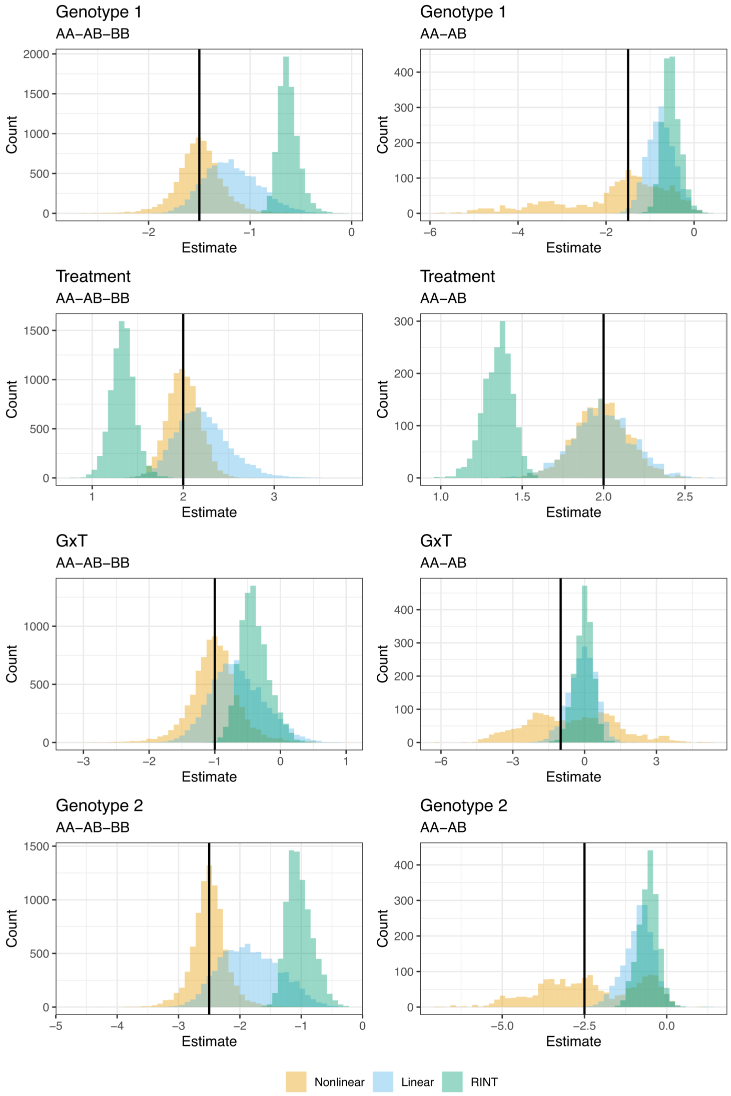

Figure S12: The same as Fig. S8 but for unpaired data.

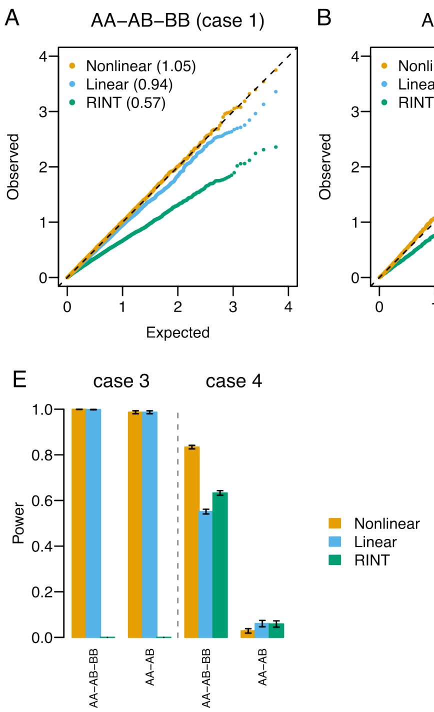

Figure S13: (A–D) Assessing null distributions in paired data. Q-Q plots comparing the theoretical null $P$ values and empirical values obtained by different methods. The values are negative $\log_{10}$-transformed. Shown in parentheses are genomic inflation factors [28]. (E) Assessing the power to detect G×T interactions in paired data. Shown is statistical power for different methods. The vertical bars represent standard deviations.

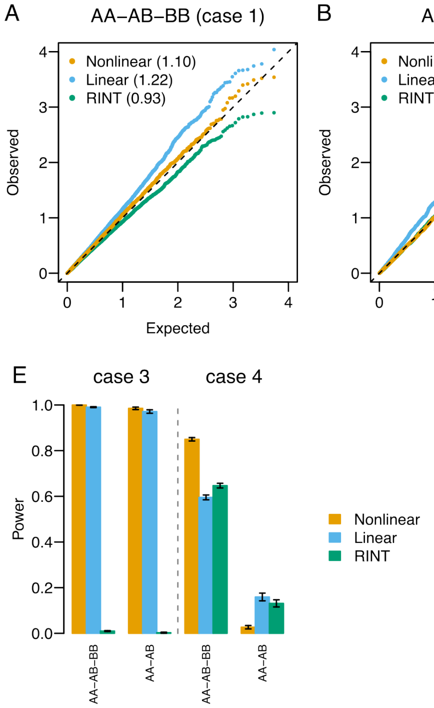

Figure S14: The same as Fig. S13 but for unpaired data.

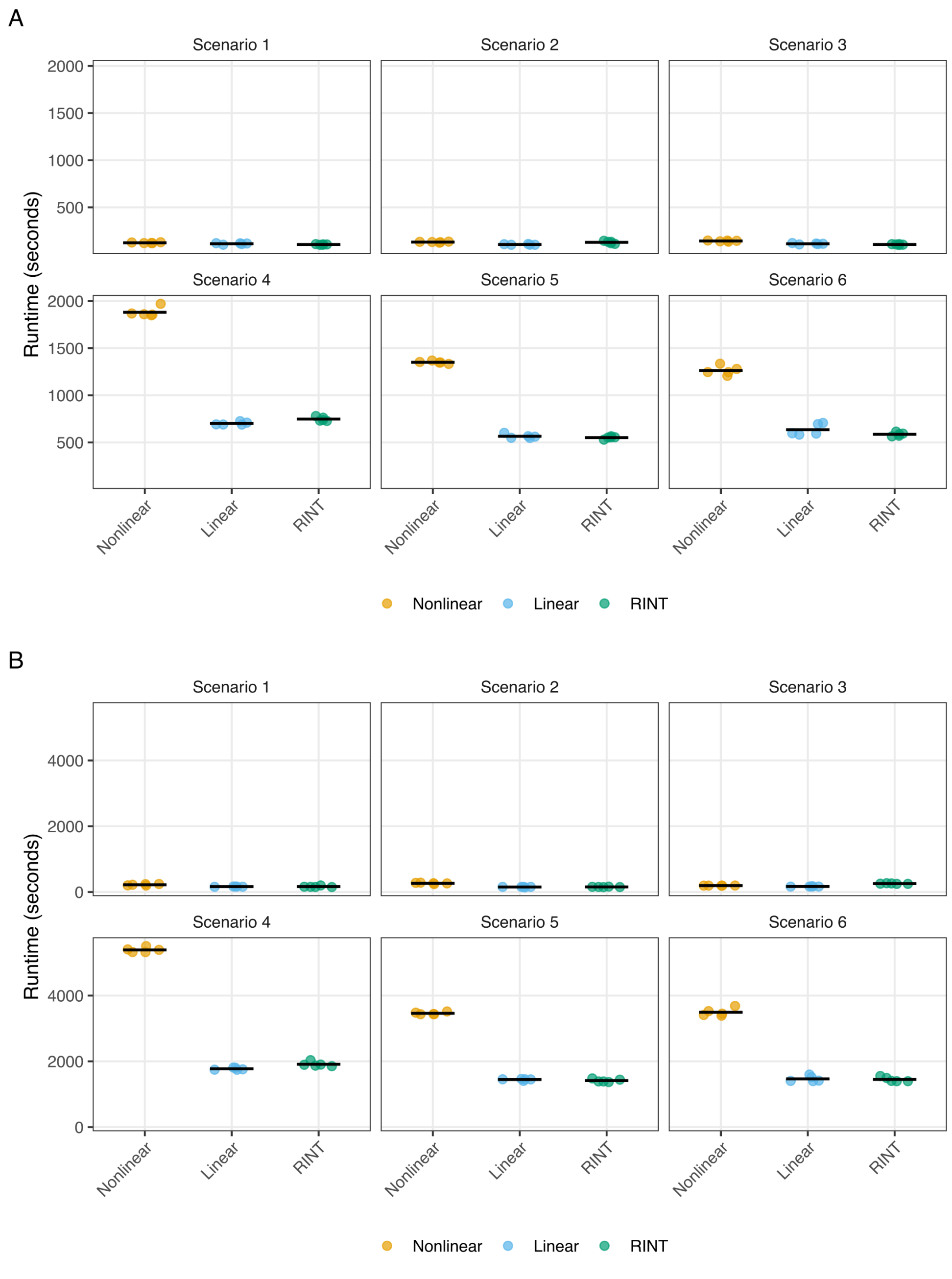

Figure S15: The runtime for performing interaction molecular QTL mapping for 10,000 paired data in six scenarios. In (A) and (B), sample sizes were 156 and 312, respectively. Points represent five repeated measurements. The horizontal bars represent mean values.

A

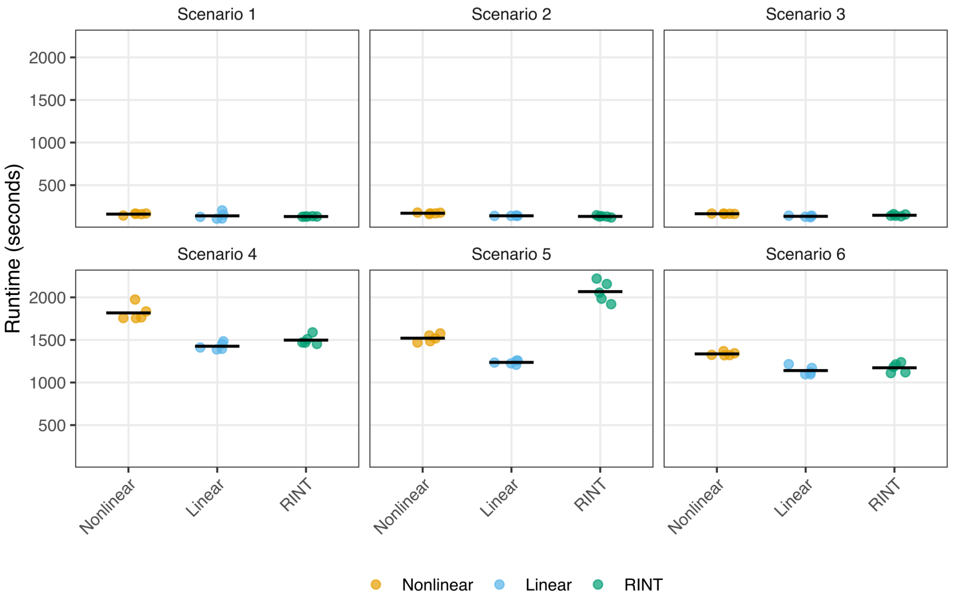

B

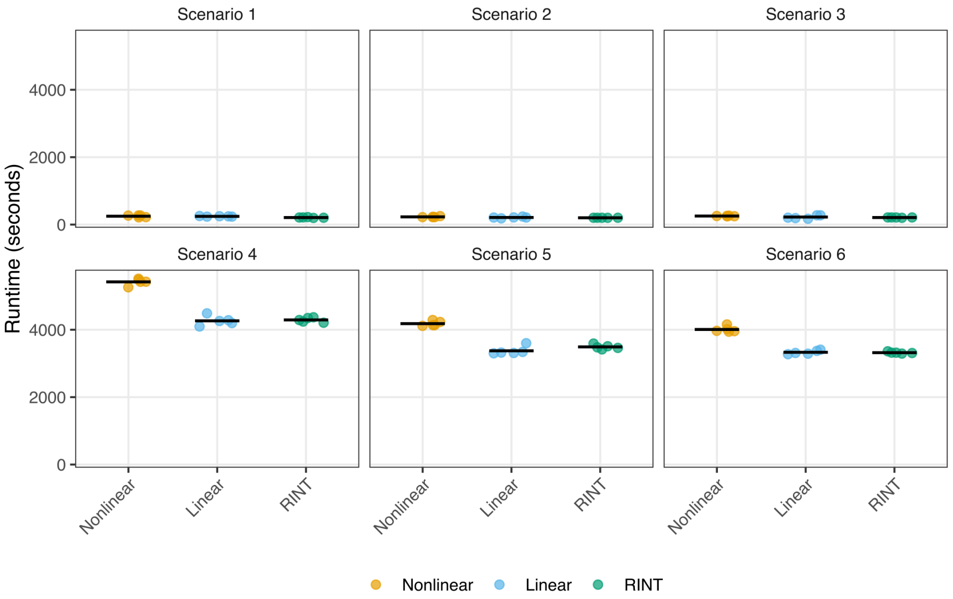

Figure S16: Same as Fig. S15 but for unpaired data.

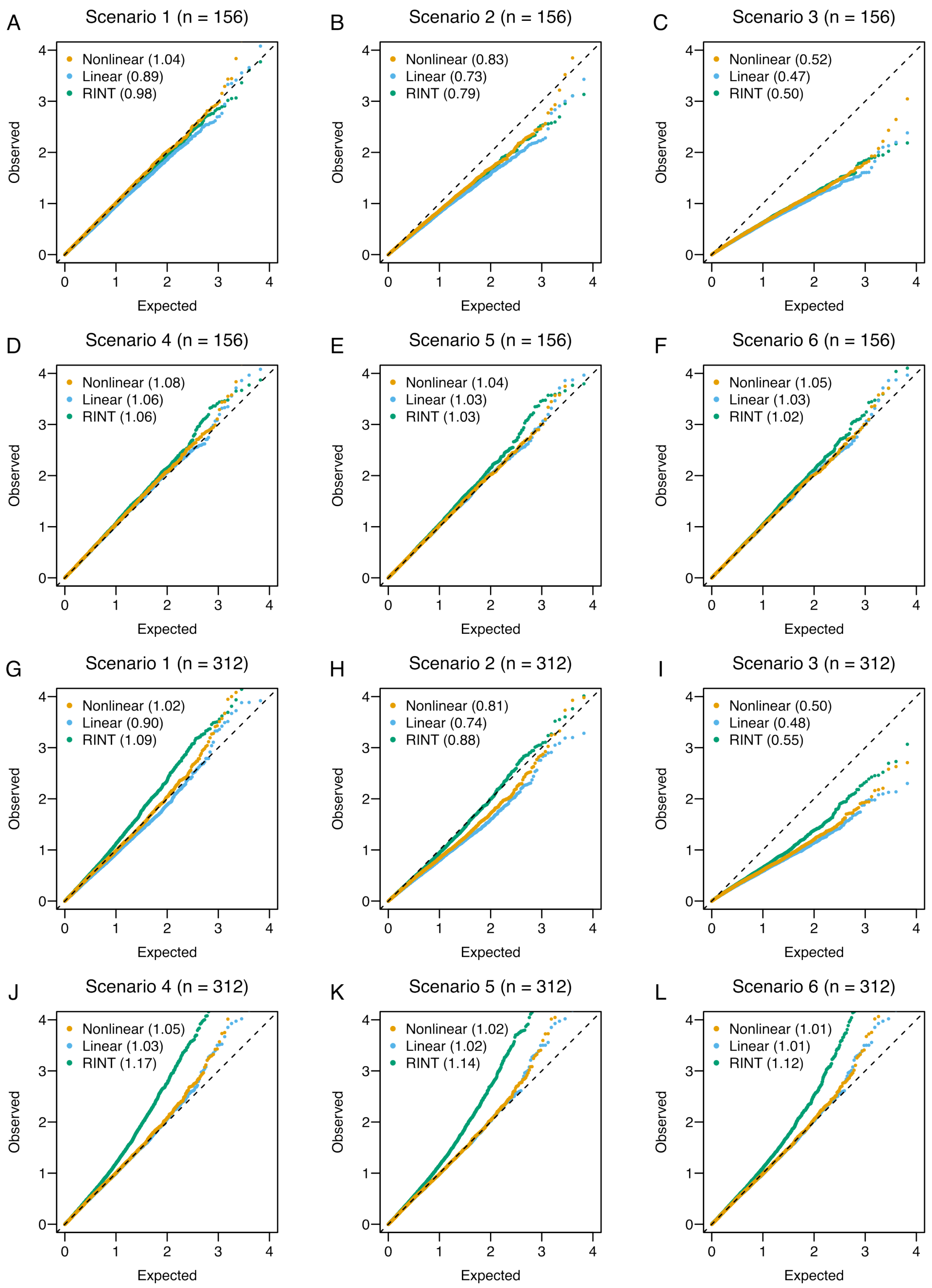

Figure S17: Assessing null distributions in paired data. Q-Q plots comparing the theoretical null $P$ values and empirical values obtained by different methods. The values are negative $\log_{10}$-transformed. Shown in parentheses are genomic inflation factors [28].

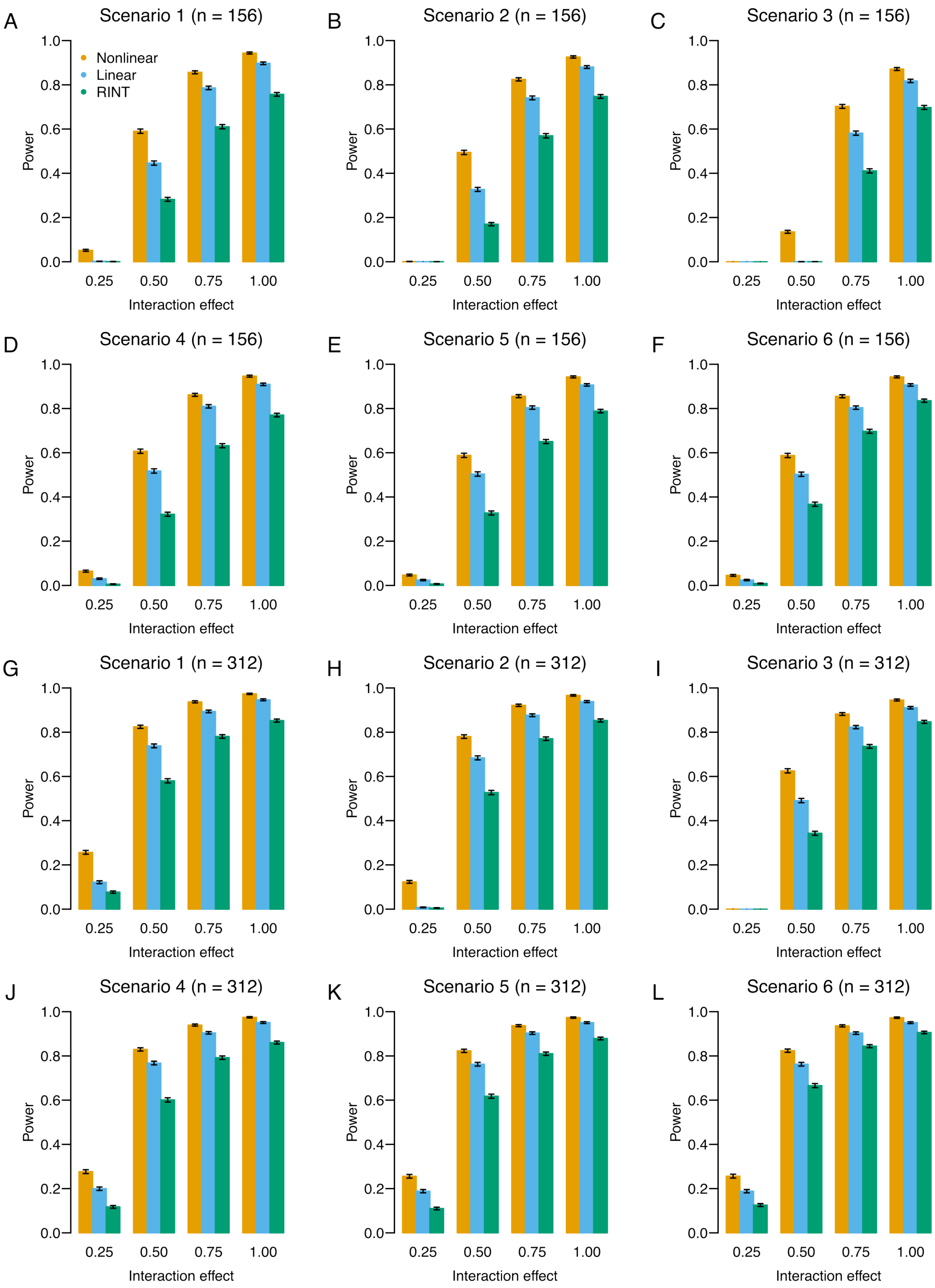

Figure S18: Assessing the power to detect G×T interactions in paired data. Shown is statistical power at varying magnitudes of G×T interaction for different methods. The vertical bars represent standard deviations.

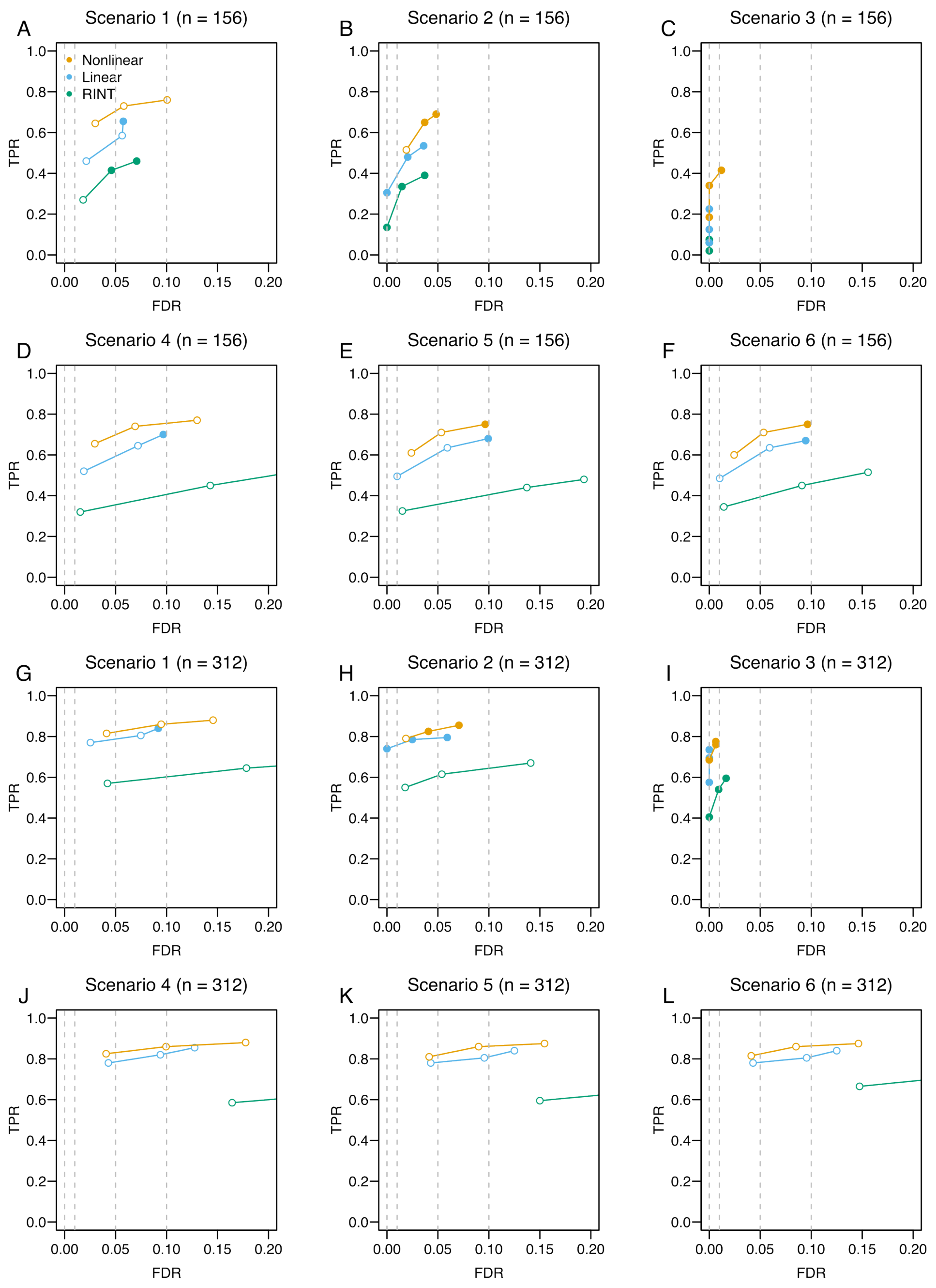

Figure S19: Assessing the overall performance in paired data. TPR is plotted against FDR at varying thresholds for different methods. The closed circle indicates that the empirical FDR is smaller than the nominal FDR. The open circle indicates otherwise.

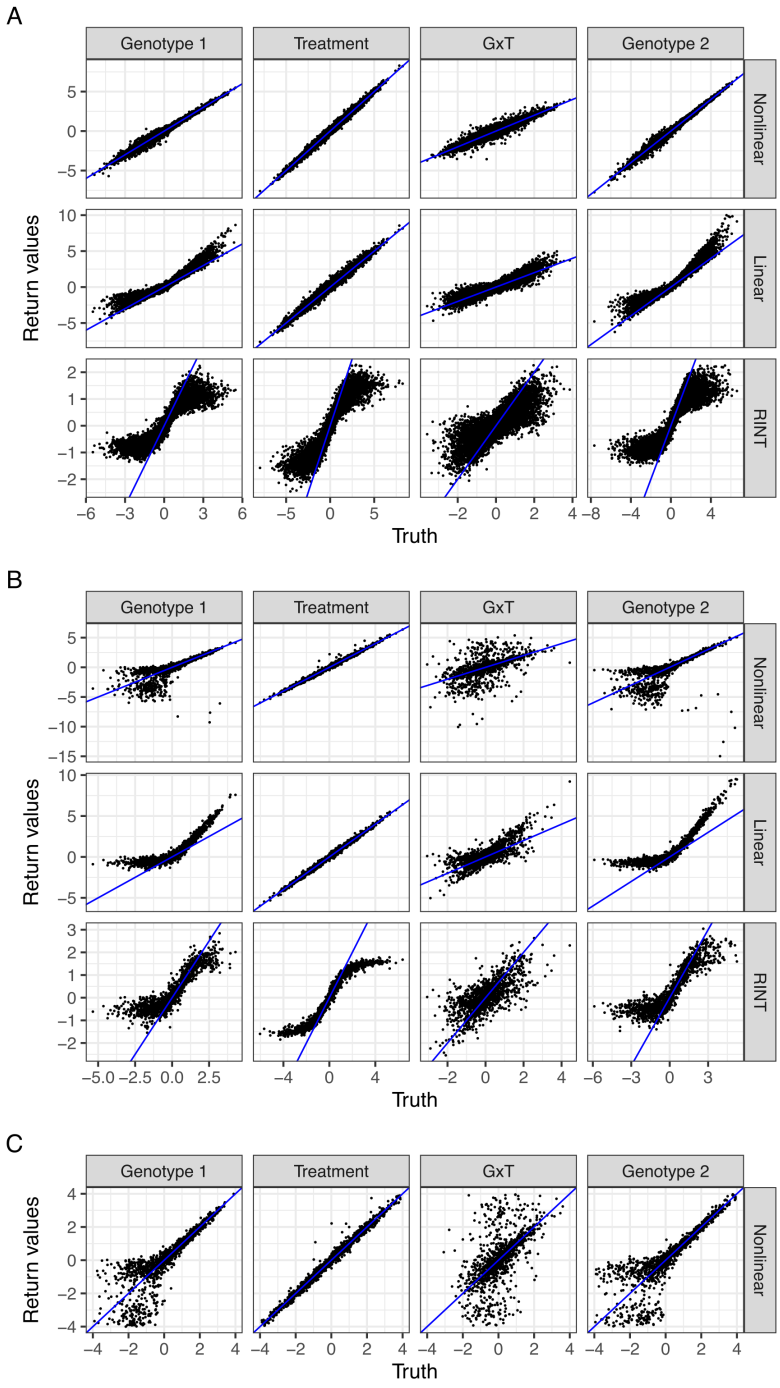

Figure S20: Scatter plots comparing the optimization return values against the true parameter values for scenario 1 and paired data. The labels Genotype 1, Treatment, G×T, and Genotype 2 represent genotype effects in condition 1, treatment effects, G×T interaction effects, and genotype effects in condition 2, respectively. (A) SNPs with three genotype levels. (B) AA-AB SNPs. (C) Zoom-in of nonlinear regression results for AA-AB SNPs.

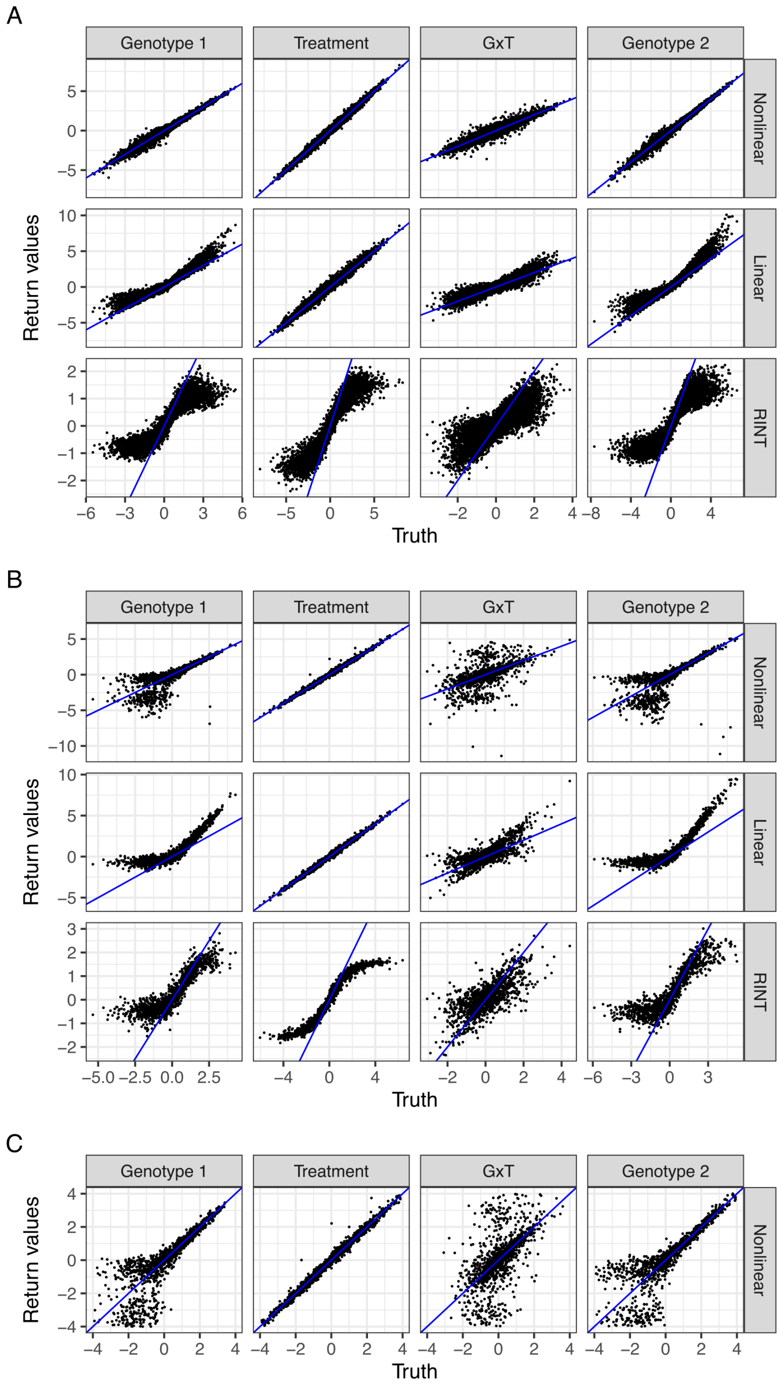

Figure S21: Same as Fig. S20 but for scenario 2.

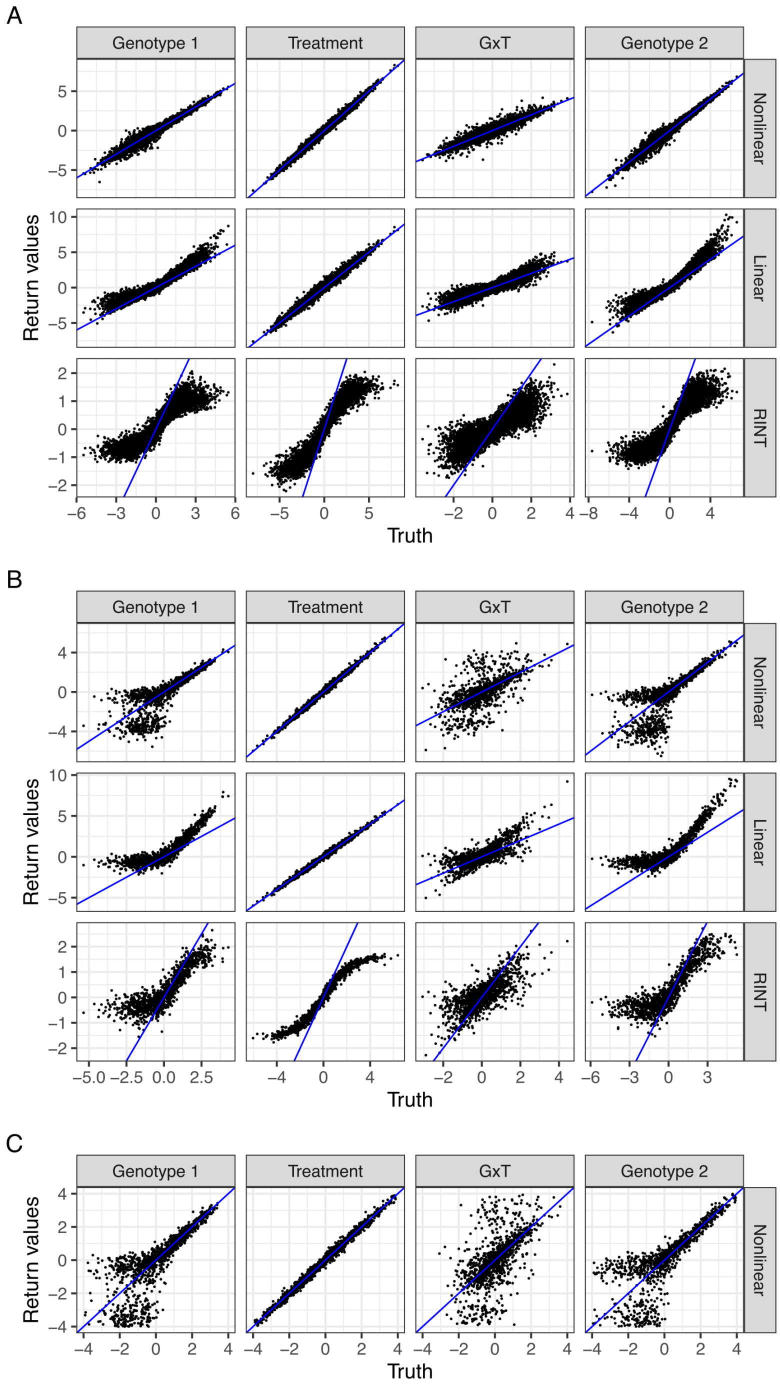

Figure S22: Same as Fig. S20 but for scenario 3.

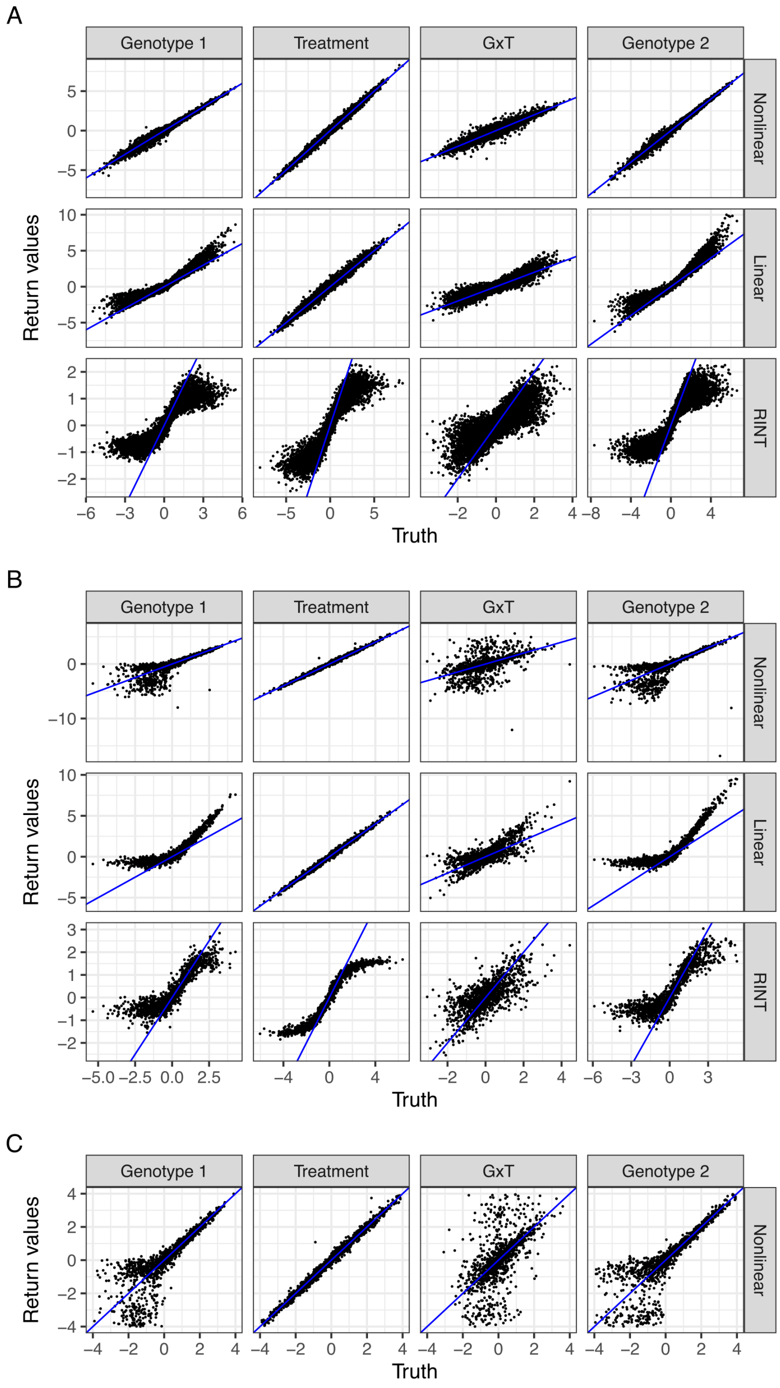

Figure S23: Same as Fig. S20 but for scenario 4.

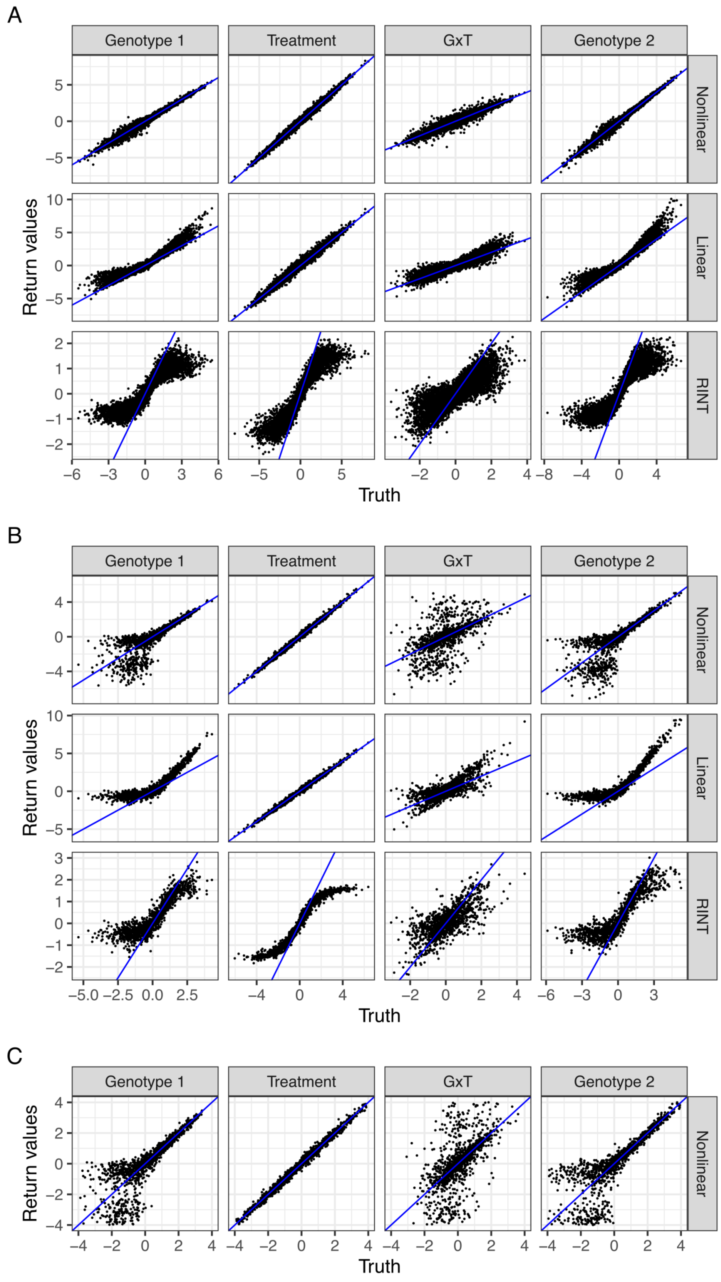

Figure S24: Same as Fig. S20 but for scenario 5.

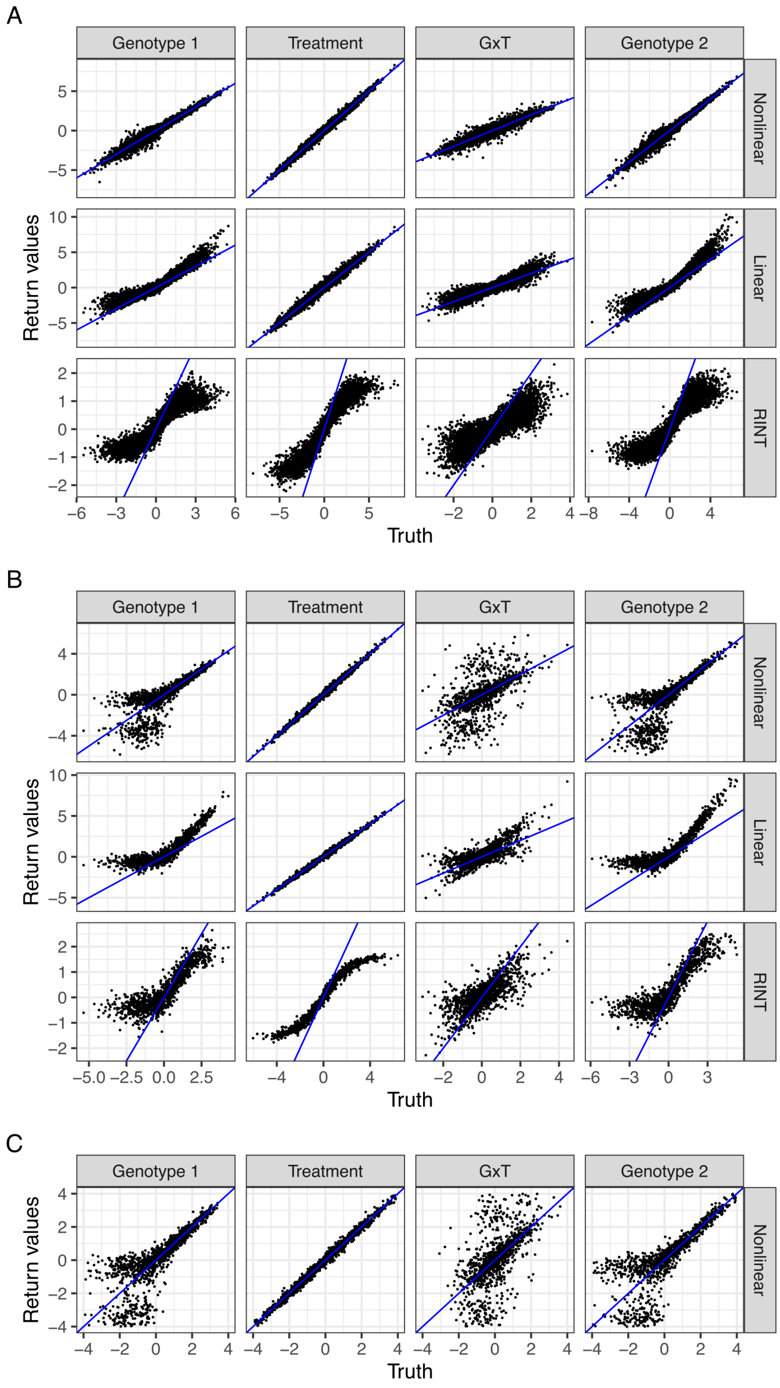

Figure S25: Same as Fig. S20 but for scenario 6.

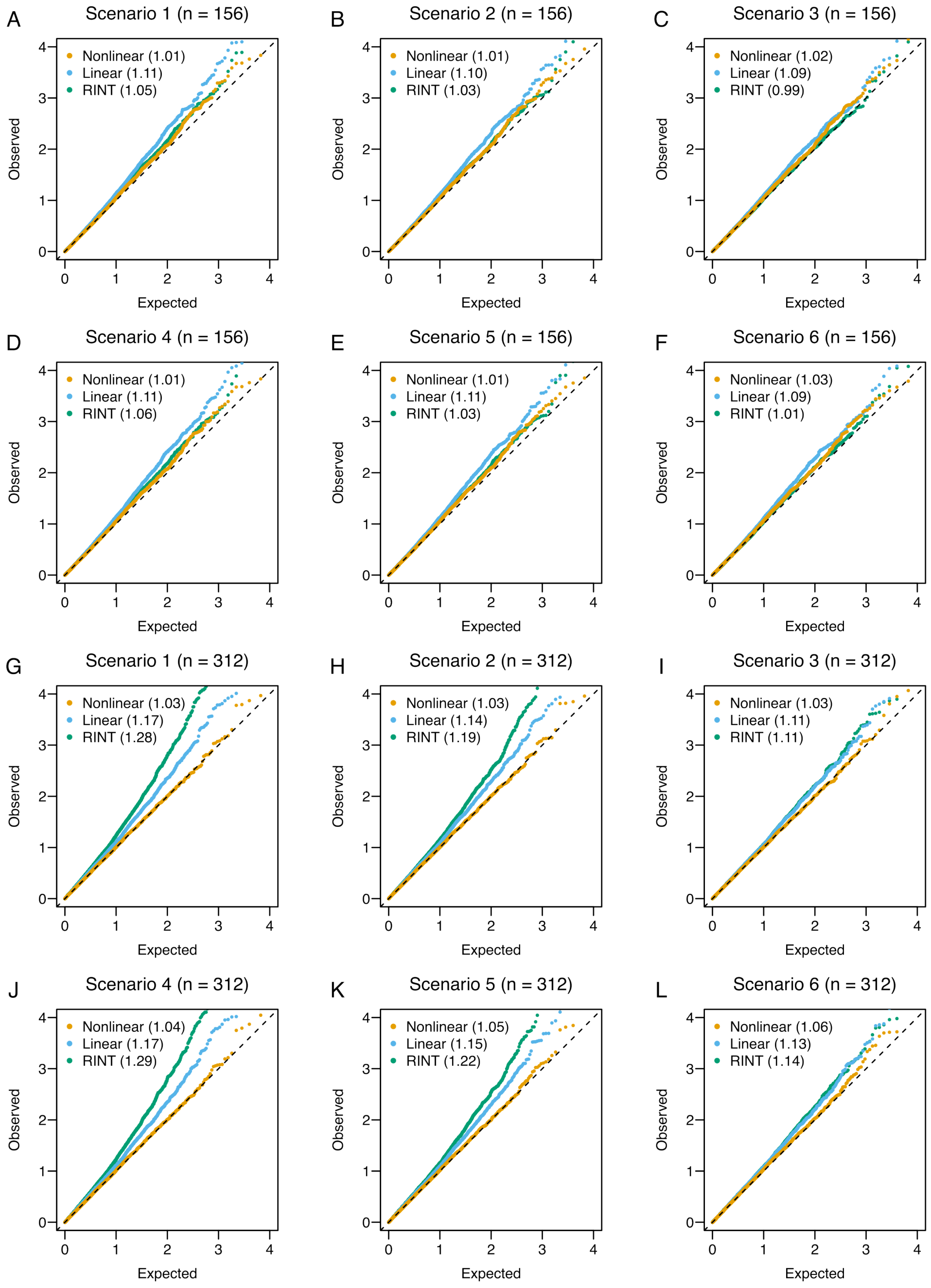

Figure S26: Assessing null distributions in unpaired data using a GRM from AFR samples of the 1000 Genomes Project. Q-Q plots comparing the theoretical null $P$ values and empirical values obtained by different methods. The values are negative $\log_{10}$-transformed. Shown in parentheses are genomic inflation factors [28].

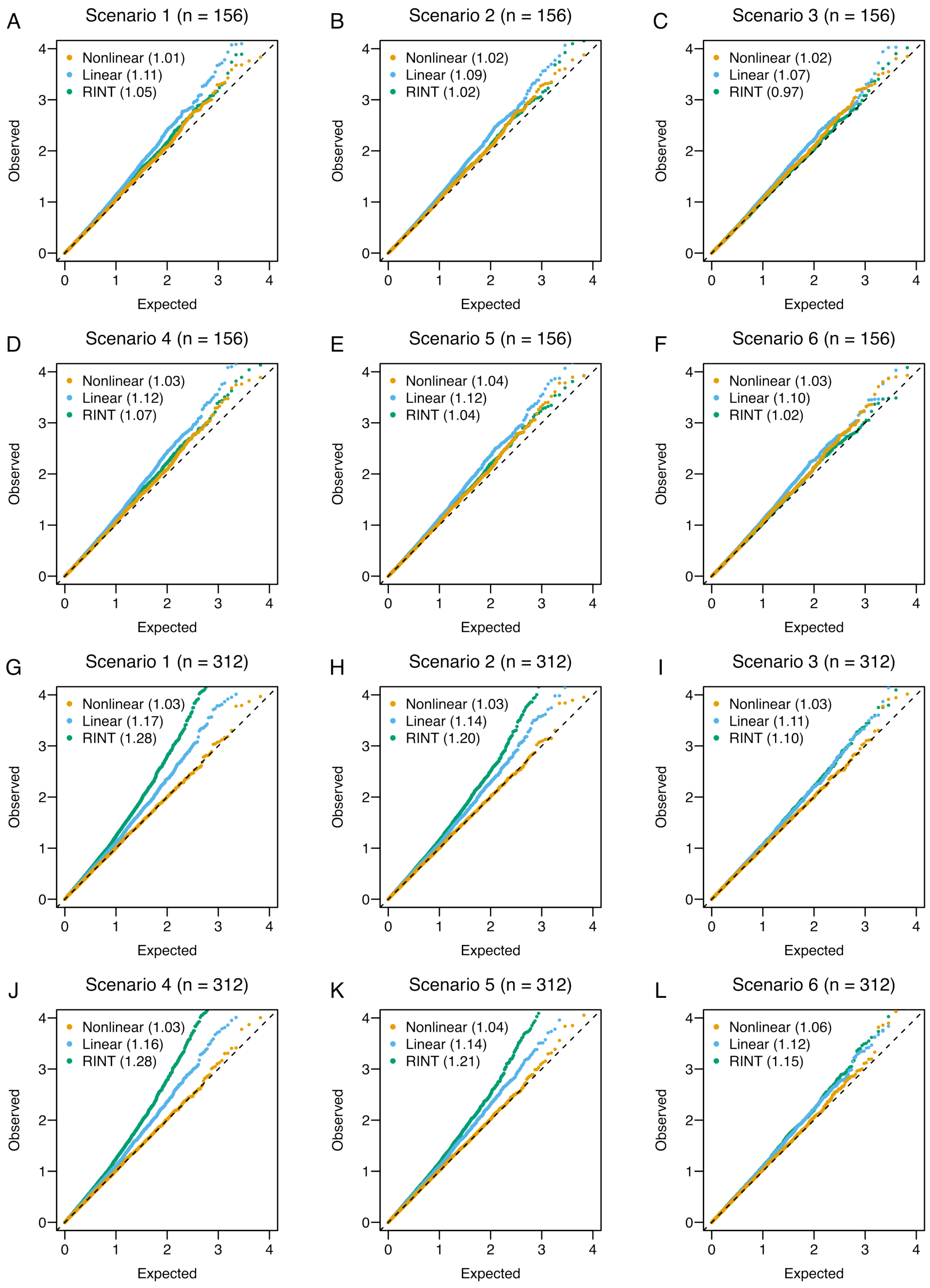

Figure S27: Same as Fig. S26 but using a GRM from EUR samples of the 1000 Genomes Project.

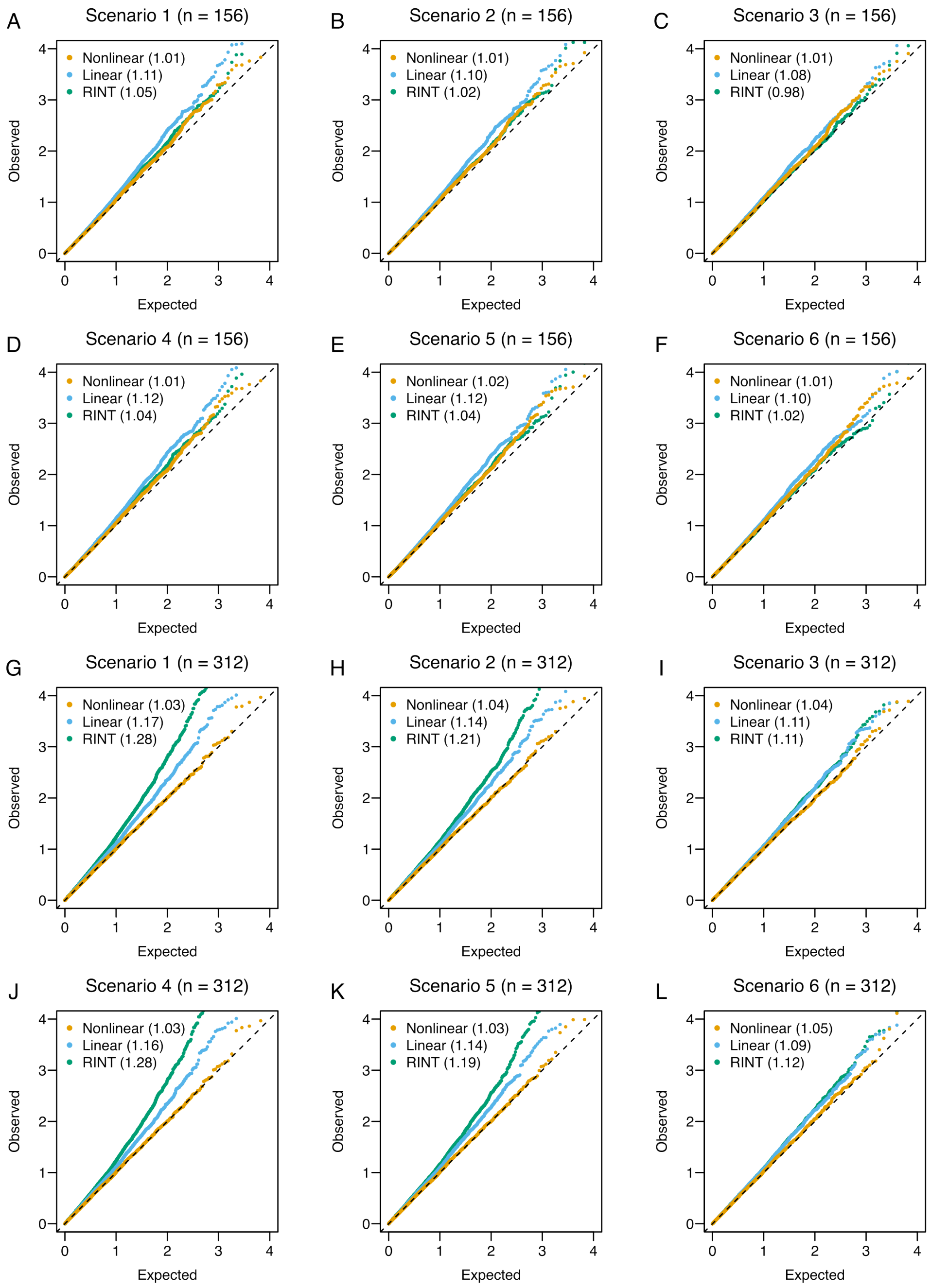

Figure S28: Same as Fig. S26 but using a GRM from EAS samples of the 1000 Genomes Project.

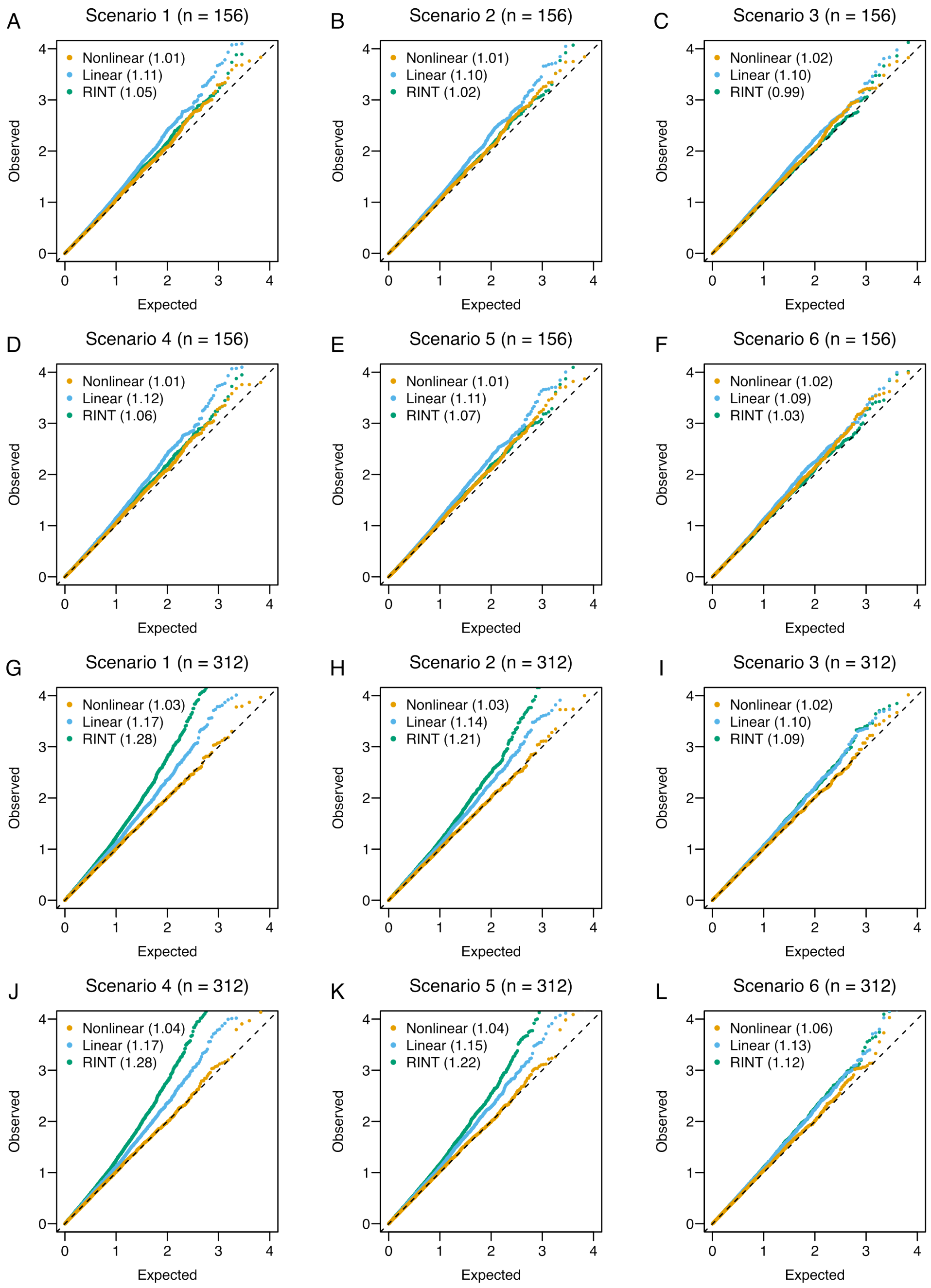

Figure S29: Same as Fig. S26 but using a GRM from SAS samples of the 1000 Genomes Project.

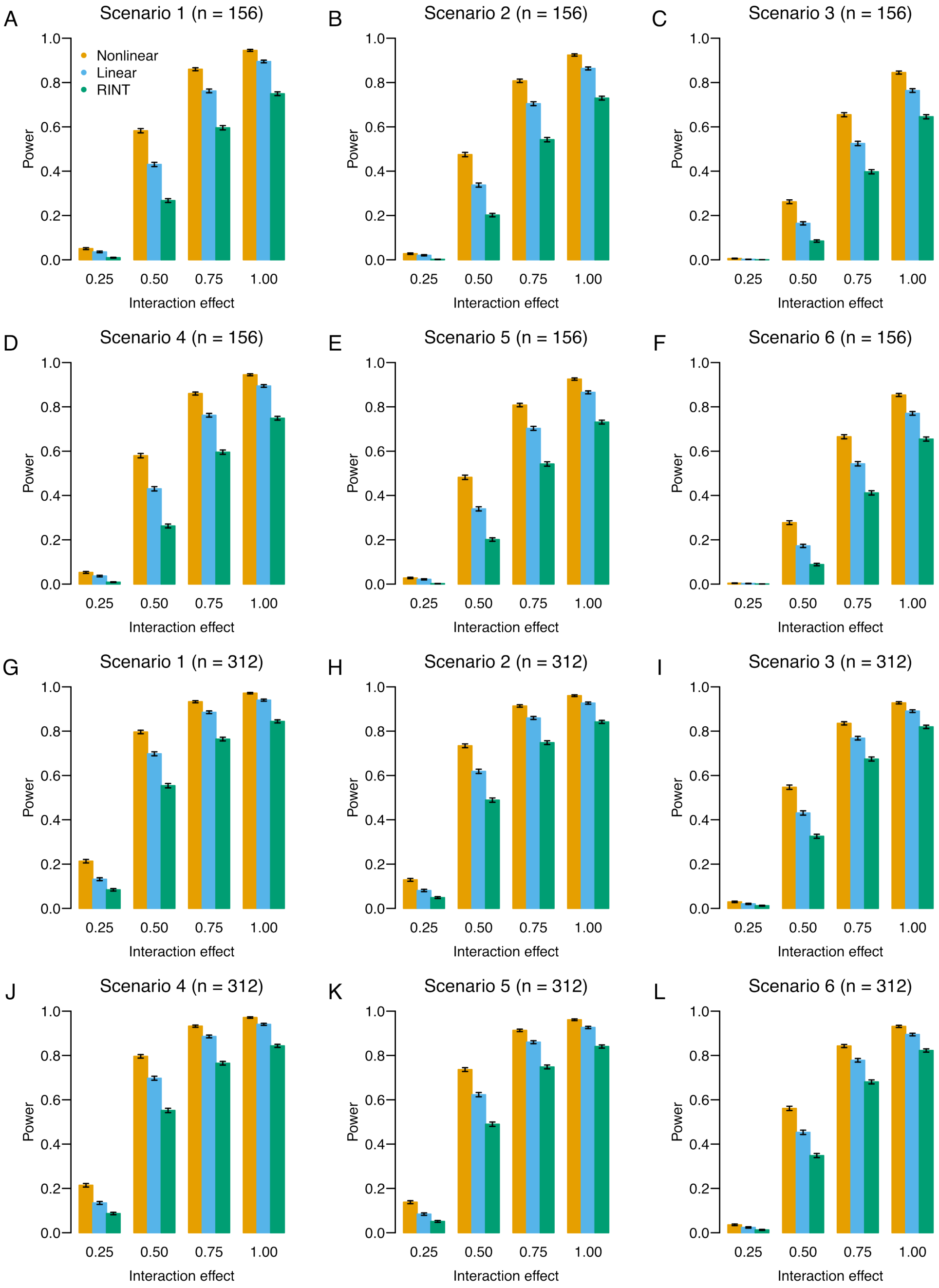

Figure S30: Assessing the power to detect G×T interactions in unpaired data using a GRM from AFR samples of the 1000 Genomes Project. Shown is statistical power at varying magnitudes of G×T interaction for different methods. The vertical bars represent standard deviations.

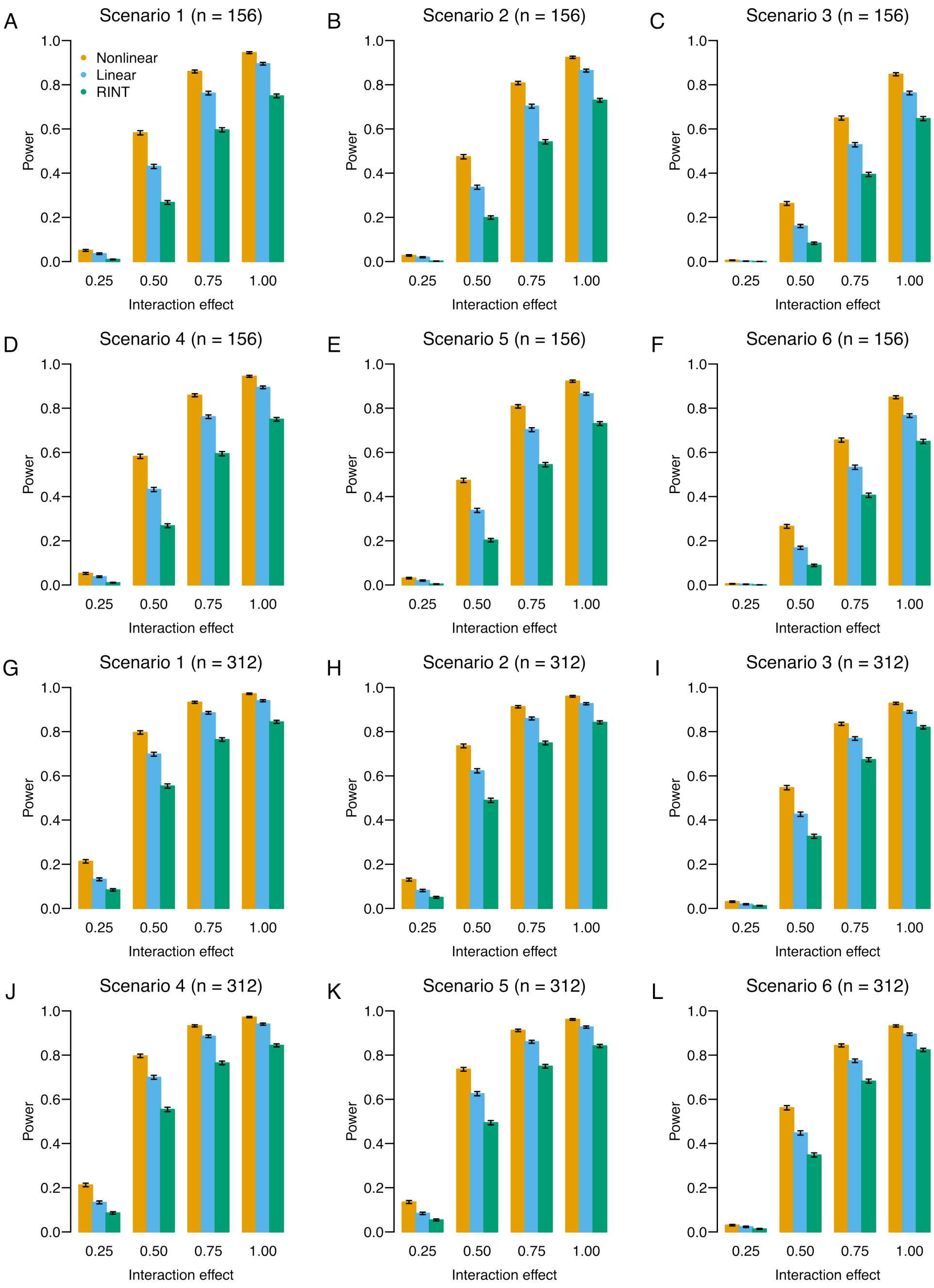

Figure S31: Same as Fig. S30 but using a GRM from EUR samples of the 1000 Genomes Project.

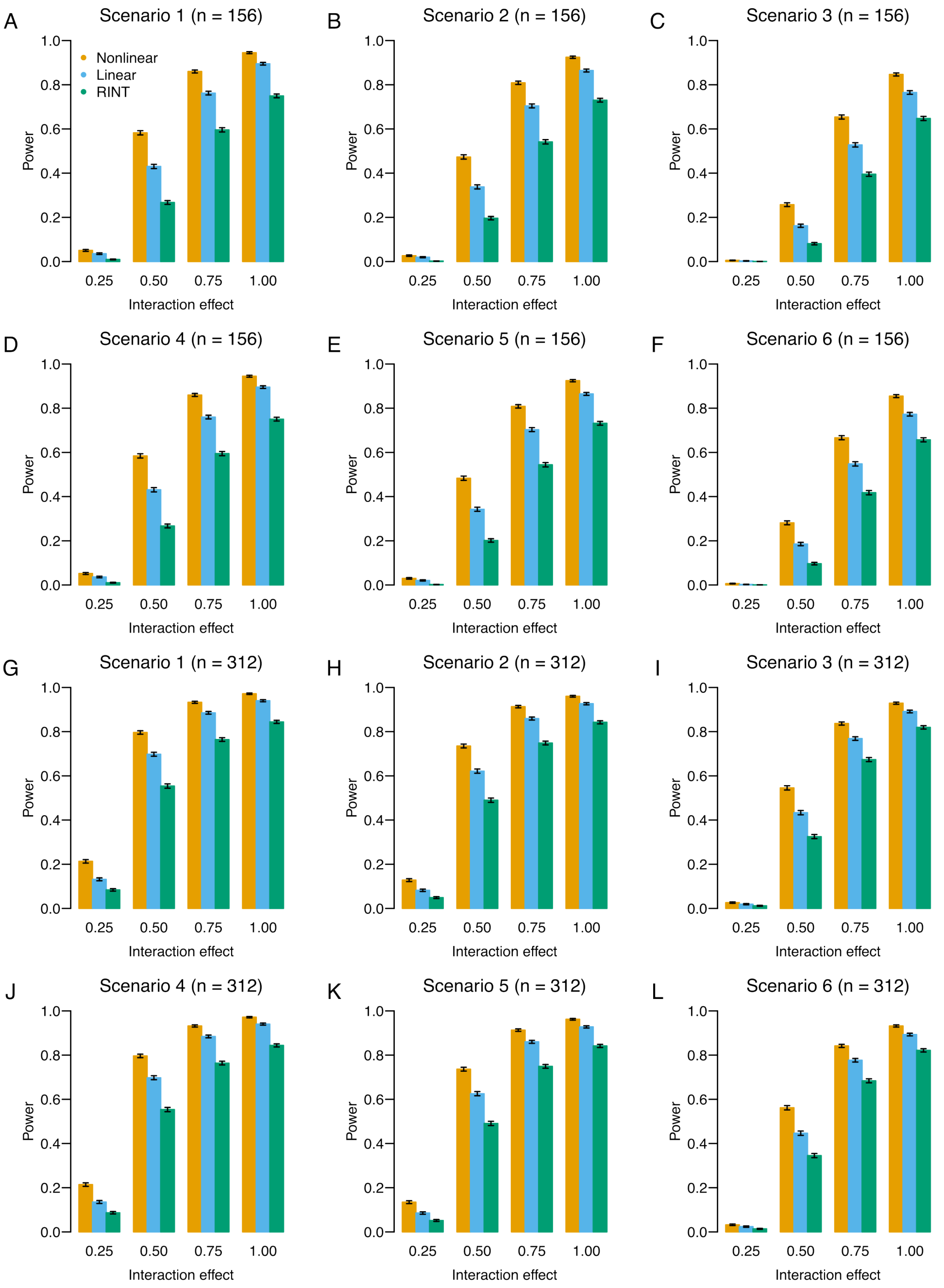

Figure S32: Same as Fig. S30 but using a GRM from EAS samples of the 1000 Genomes Project.

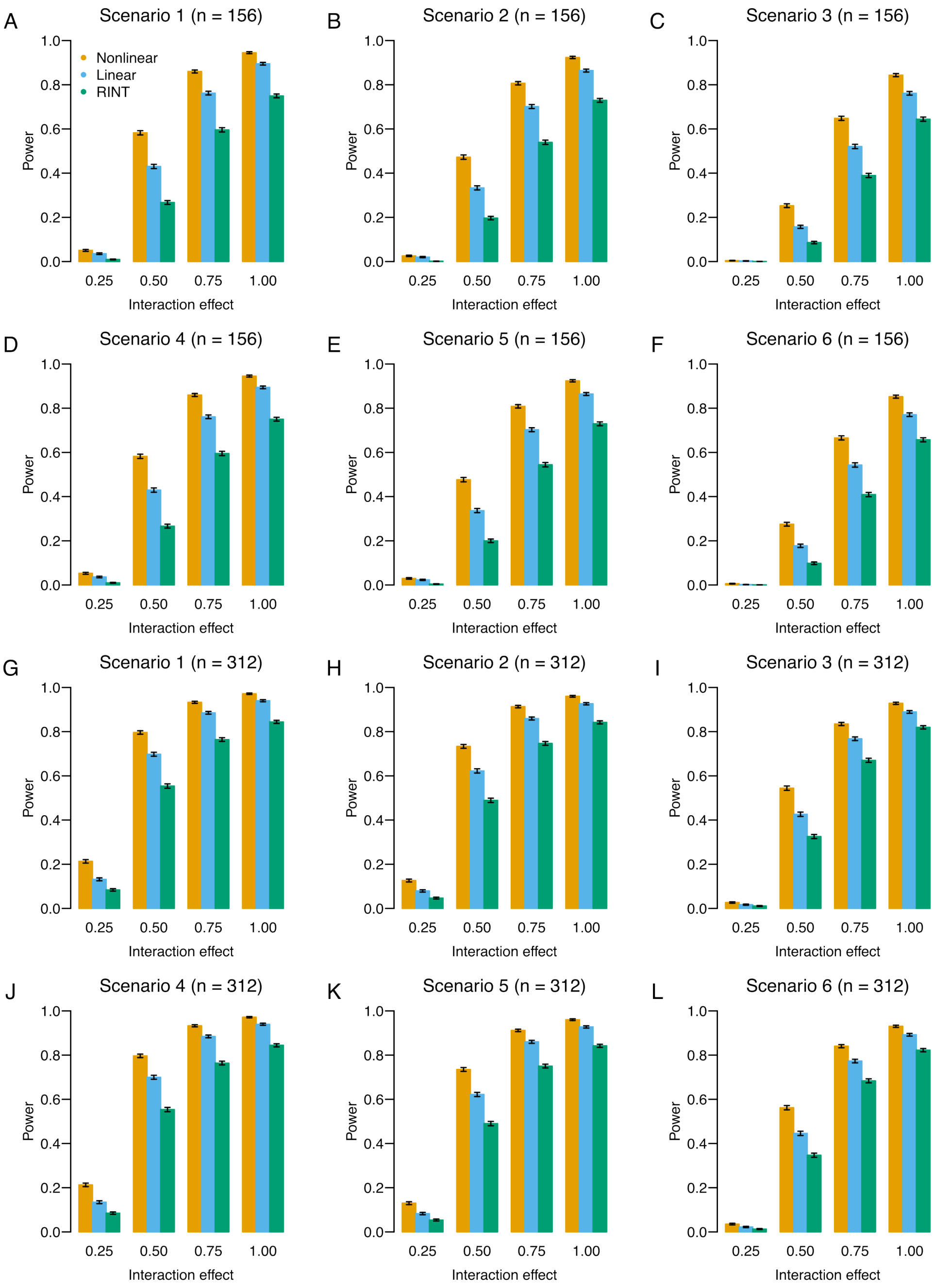

Figure S33: Same as Fig. S30 but using a GRM from SAS samples of the 1000 Genomes Project.

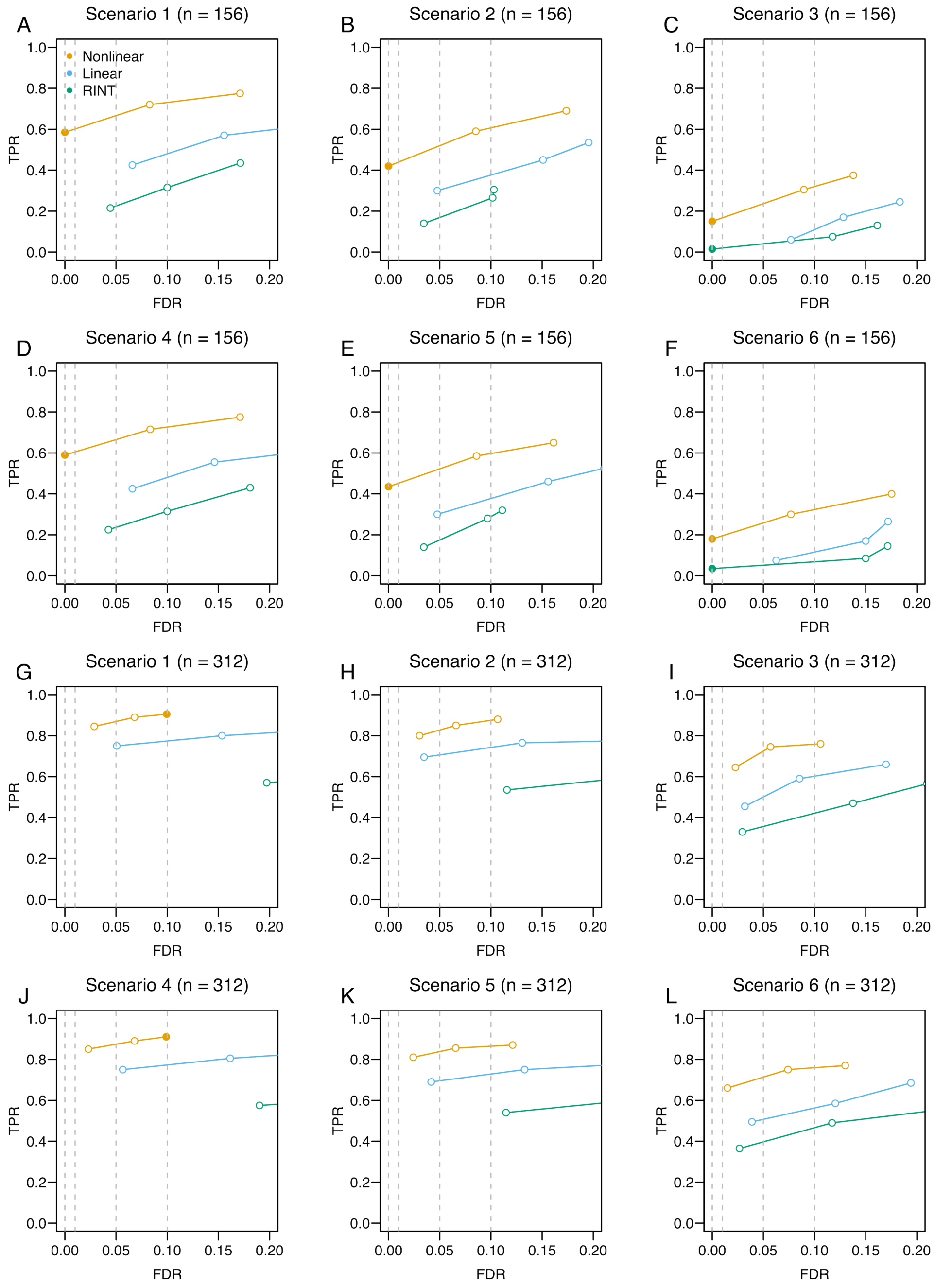

Figure S34: Assessing the overall performance in unpaired data using a GRM from AFR samples of the 1000 Genomes Project. TPR is plotted against FDR at varying thresholds for different methods. The closed circle indicates that the empirical FDR is smaller than the nominal FDR. The open circle indicates otherwise.

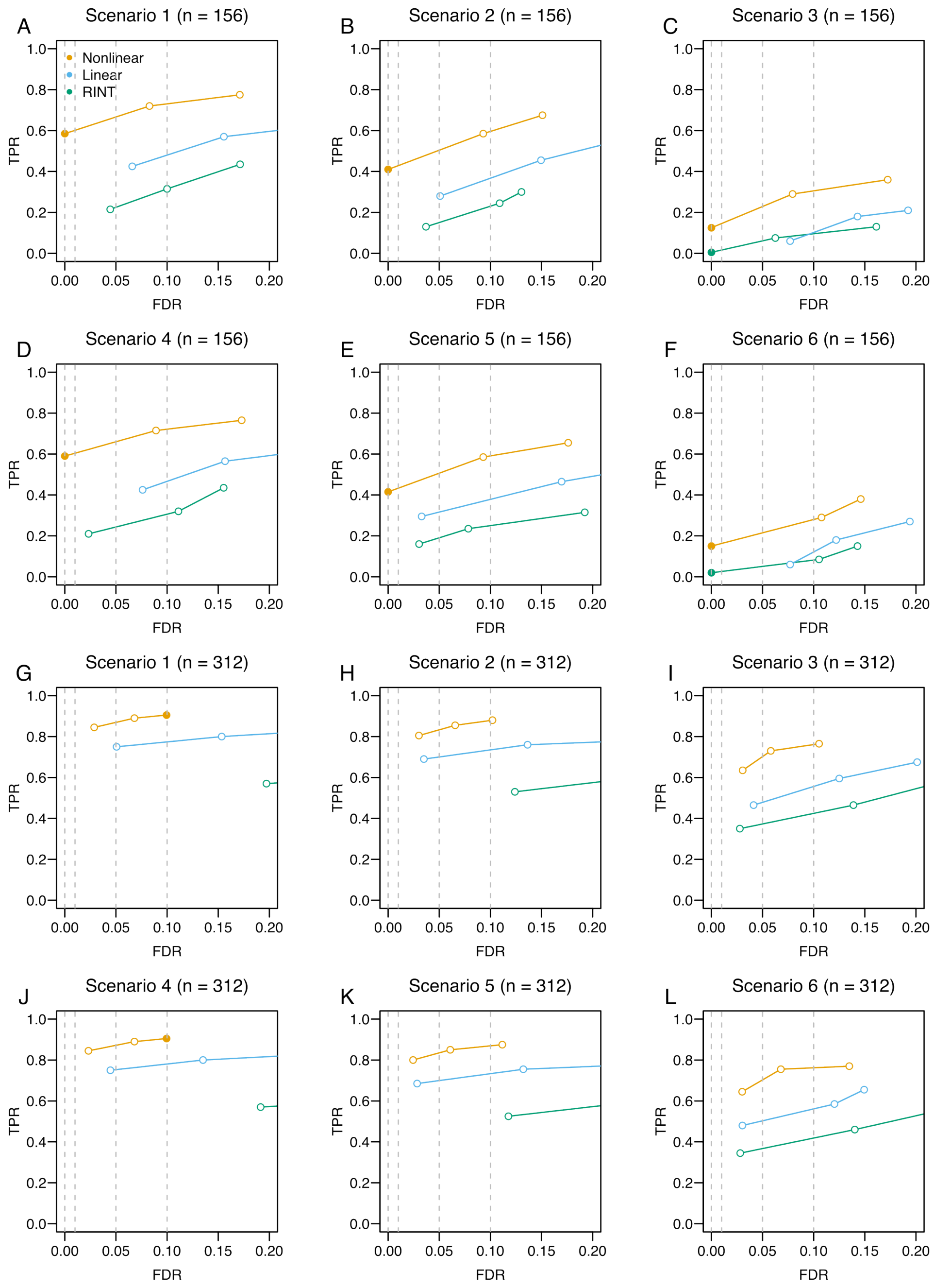

Figure S35: Same as Fig. S34 but using a GRM from EUR samples of the 1000 Genomes Project.

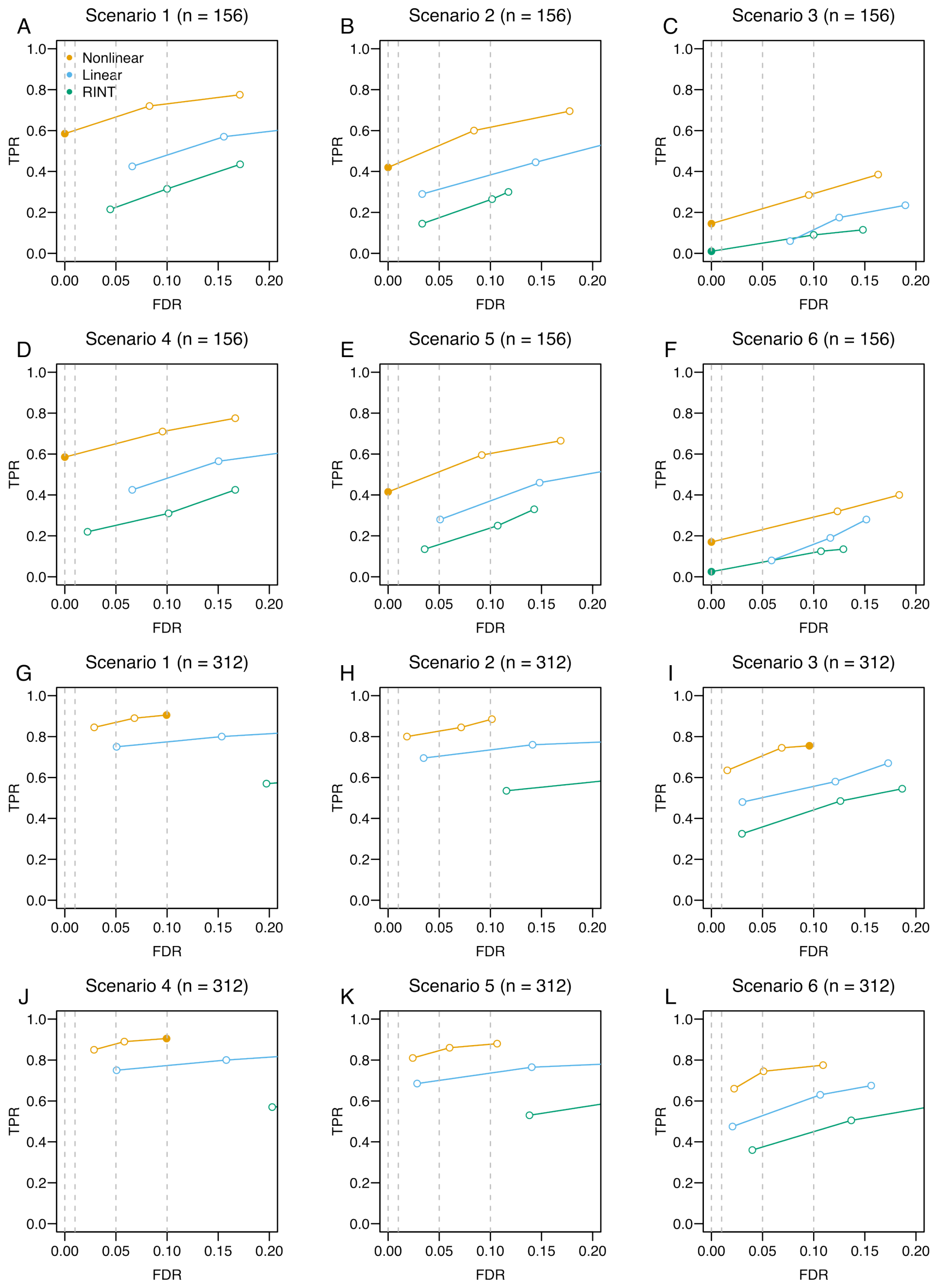

Figure S36: Same as Fig. S34 but using a GRM from EAS samples of the 1000 Genomes Project.

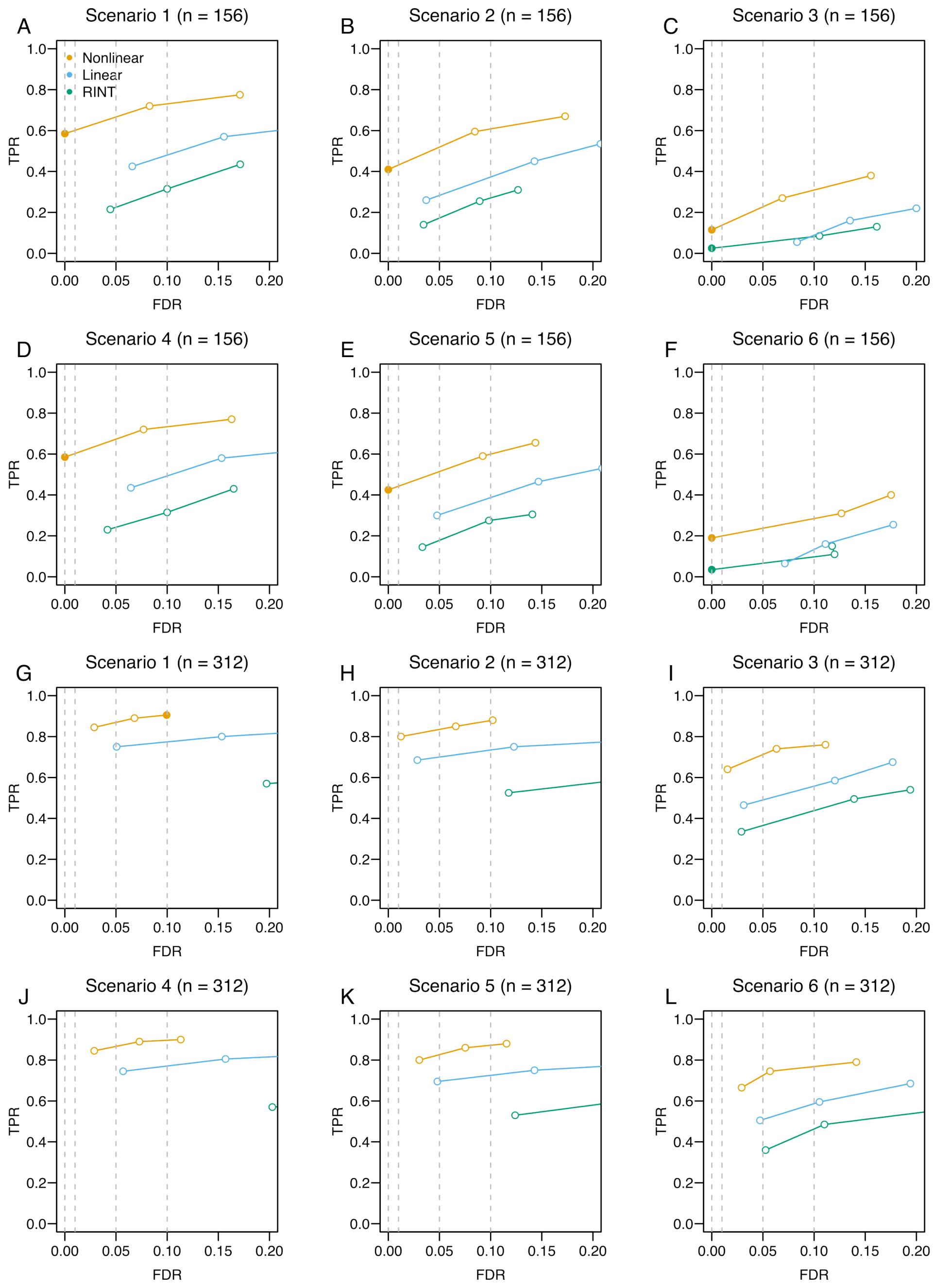

Figure S37: Same as Fig. S34 but using a GRM from SAS samples of the 1000 Genomes Project.

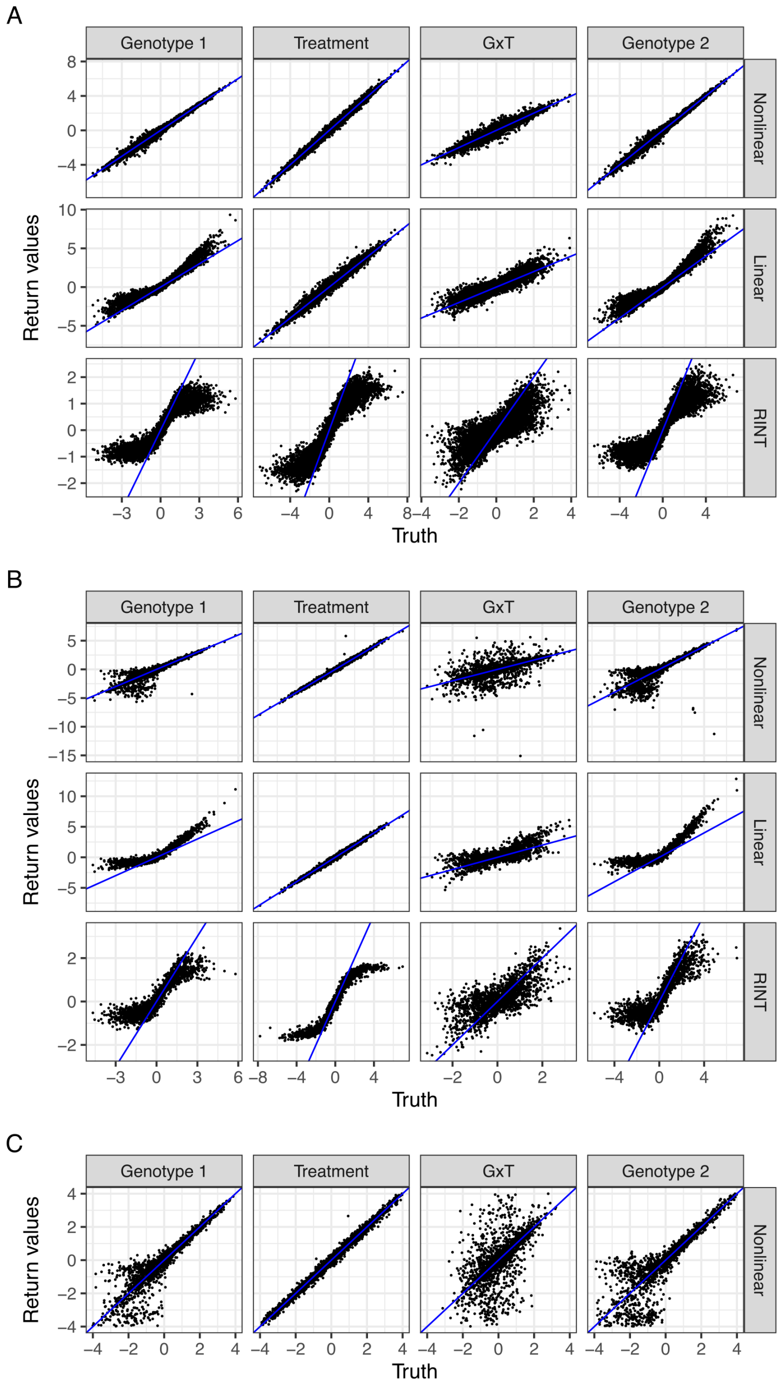

Figure S38: Same as Fig. S20 but for unpaired data.

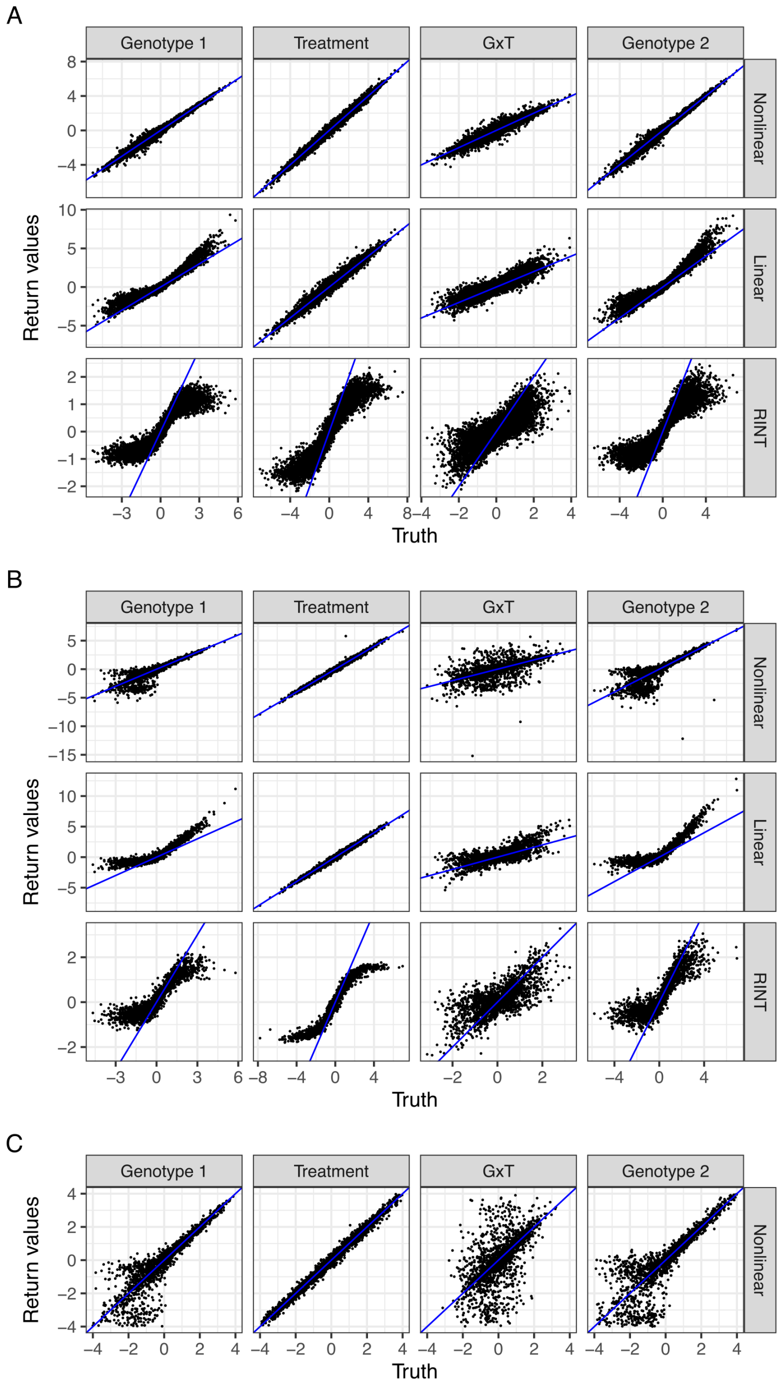

Figure S39: Same as Fig. S38 but for scenario 2.

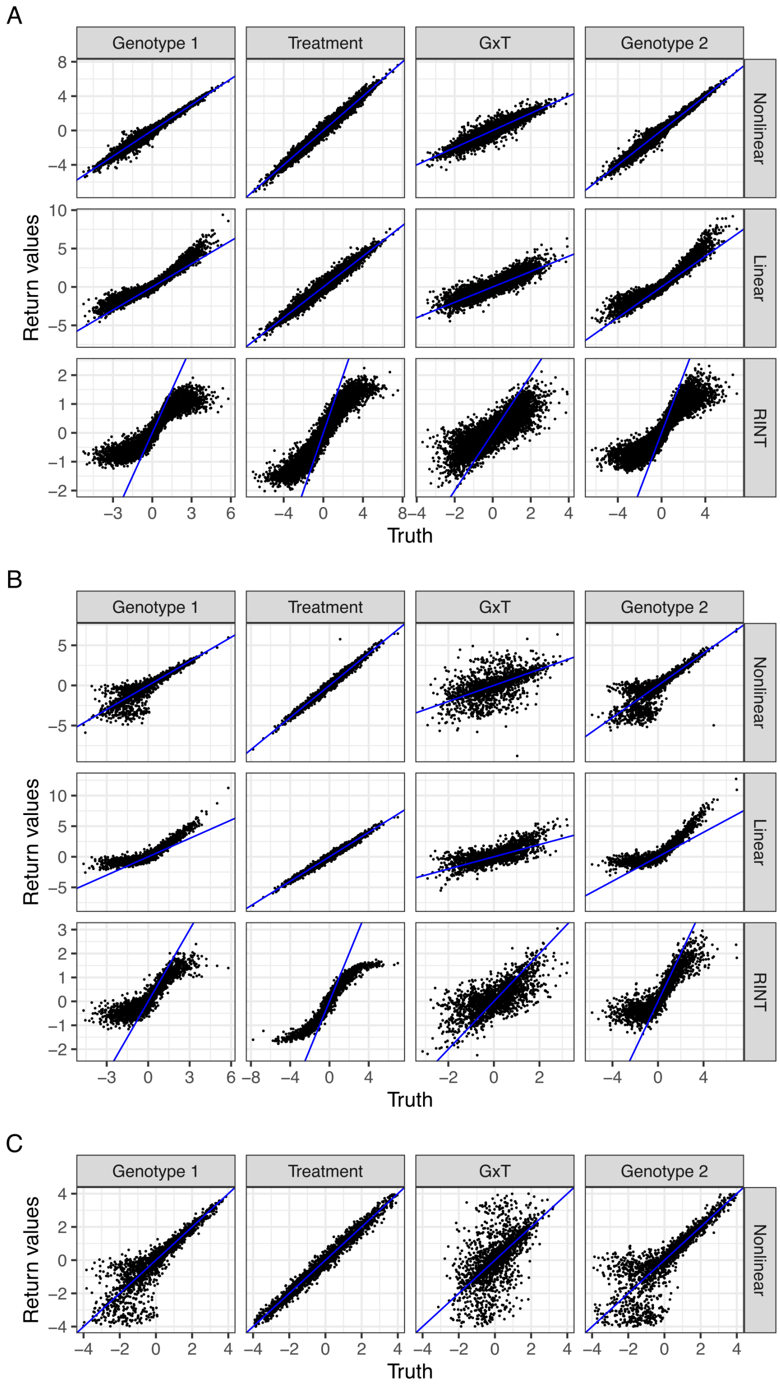

Figure S40: Same as Fig. S38 but for scenario 3.

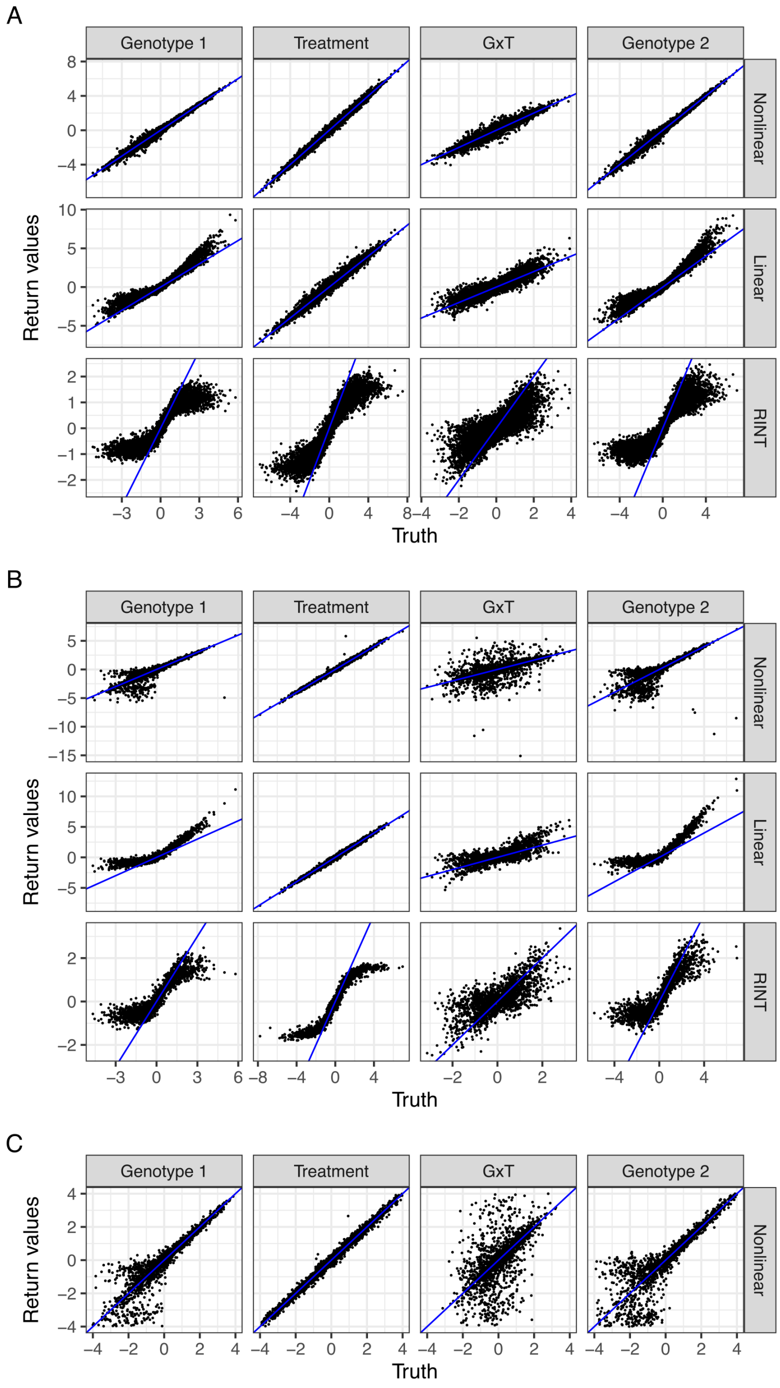

Figure S41: Same as Fig. S38 but for scenario 4.

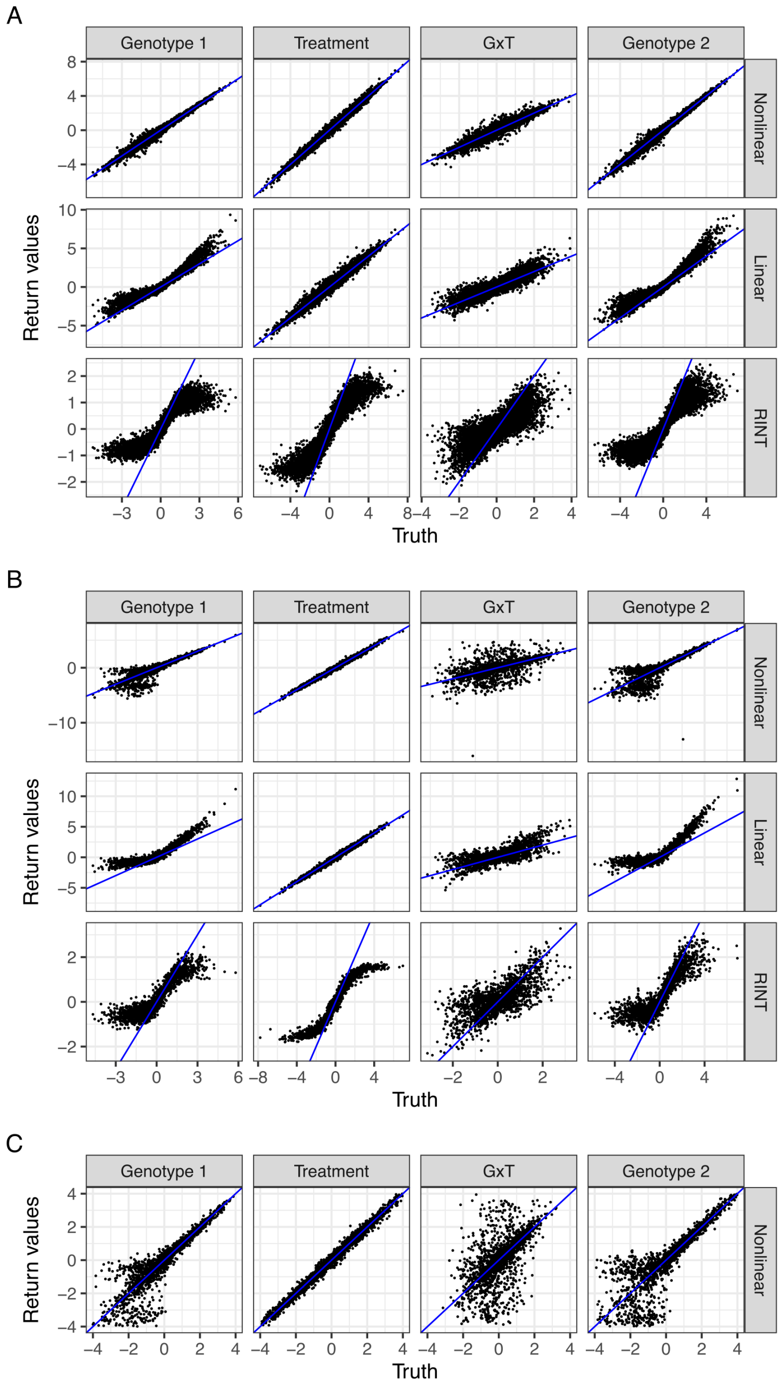

Figure S42: Same as Fig. S38 but for scenario 5.

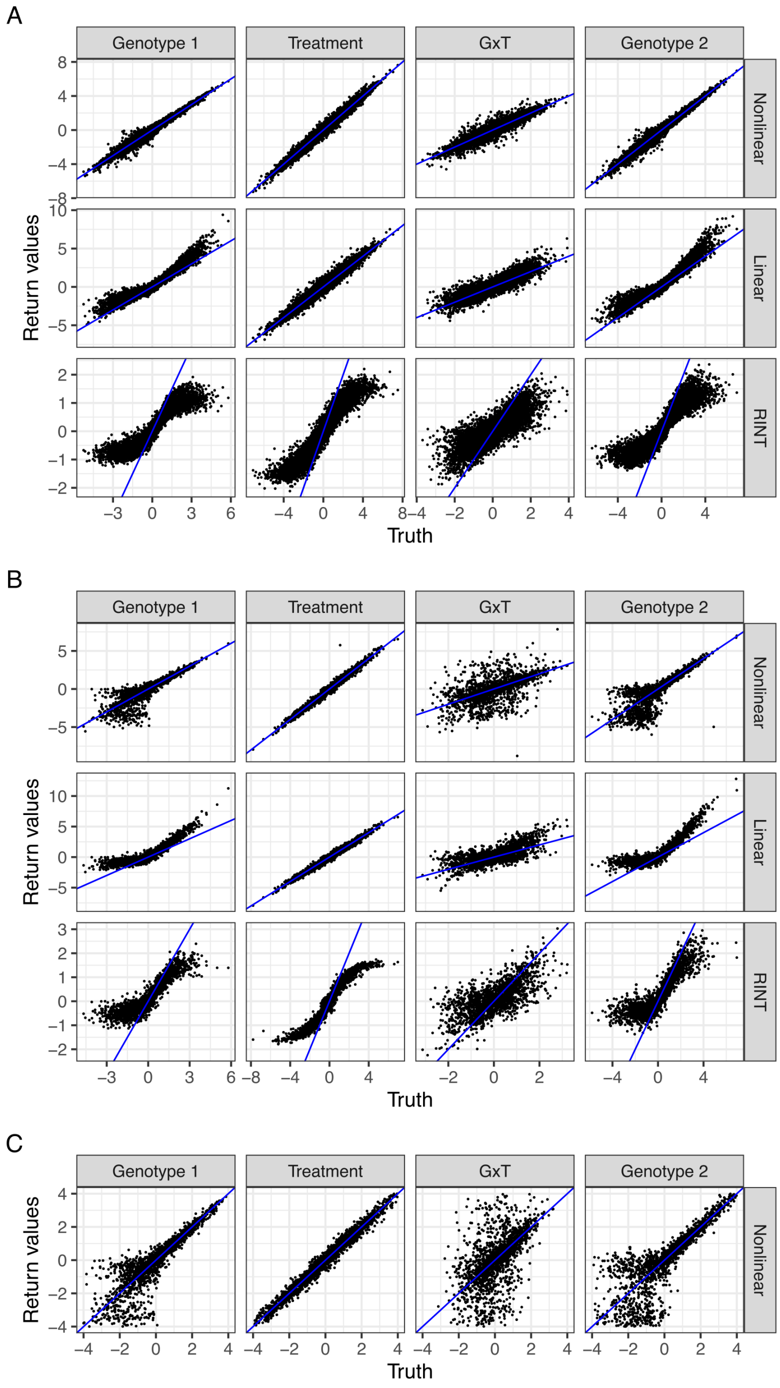

Figure S43: Same as Fig. S38 but for scenario 6.

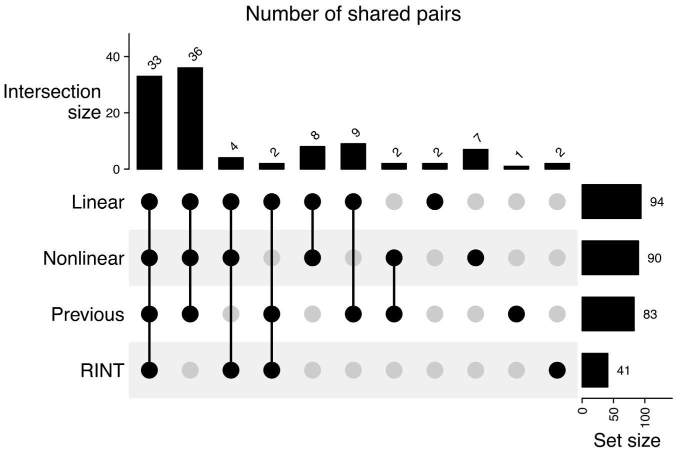

Figure S44: Overlaps between sets of gene-SNP pairs with significant G×T interactions identified by different modeling approaches. Shown is an “UpSet” plot visualizing the sizes of sets and their intersections. “Previous” corresponds to a set identified by a previous study [7].
